# Antibacterial activity of solid surfaces is critically dependent on relative humidity, inoculum volume and organic soiling

**DOI:** 10.1101/2023.03.28.534510

**Authors:** Harleen Kaur, Merilin Rosenberg, Mati Kook, Dmytro Danilian, Vambola Kisand, Angela Ivask

## Abstract

Antimicrobial surface materials potentially prevent pathogen transfer from contaminated surfaces. Efficacy of such surfaces is assessed by standard methods using wet exposure conditions known to overestimate antimicrobial activity compared to dry exposure. Some dry test formats have been proposed but semi-dry exposure scenarios *e.g.,* oral spray or water droplets exposed to ambient environment, are less studied. We aimed to determine the impact of environmental test conditions on antibacterial activity against the model species *Escherichia coli* and *Staphylococcus aureus*. Surfaces based on copper, silver, and quaternary ammonium with known or claimed antimicrobial properties were tested in conditions mimicking microdroplet spray or larger water droplets exposed to variable relative air humidity in the presence or absence of organic soiling. All the environmental parameters critically affected antibacterial activity of the tested surfaces from no effect in high-organic dry conditions to higher effect in low-organic humid conditions but not reaching the effect size demonstrated in the ISO 22169 wet format. Copper was the most efficient antibacterial surface followed by silver and quaternary ammonium based coating. Antimicrobial testing of surfaces using small droplet contamination in application-relevant conditions could therefore be considered as one of the worst-case exposure scenarios relevant to dry use surfaces.

**Featured image + One Sentence summary:** Antibacterial activity of copper and silver surfaces is highly dependent on environmental testing conditions with maximum efficiency in low-organic wet conditions to no antibacterial activity in high-organic dry conditions indicating the need to test antimicrobial surface materials in application-relevant test formats as opposed to current standards.

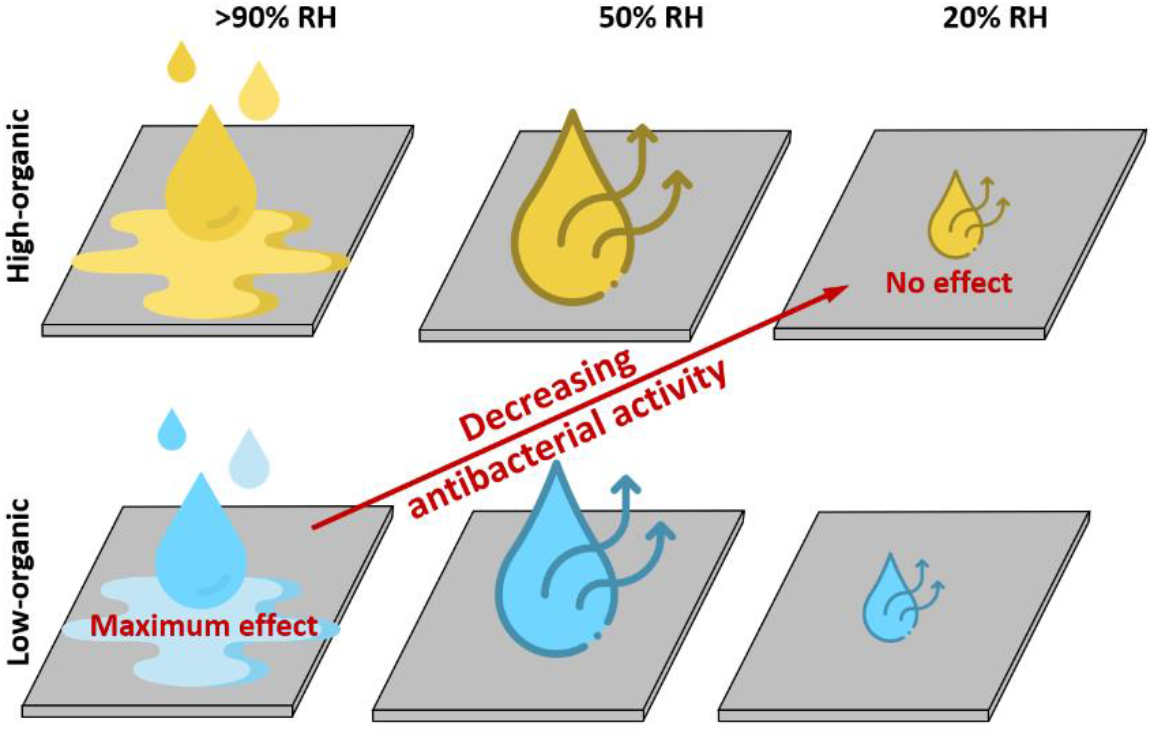

## Introduction

Surface transfer of microbes is one of the main routes of transmission of infectious diseases in critical applications including health-care environment [1]. Therefore, surfaces decreasing microbial residence time and viability and thus, the likelihood of transfer of potential pathogens, may have a significant contribution to the decrease of infectious diseases and improvement of public health [2]. However, such efficacy may only be obtained if the surface elicits sufficient antimicrobial activity. According to legislative acts and their guidance documents in the European Union [3,4] and in the United States [5,6], a proof of at least of 3 log_10_ (99.9%) reduction in microbial viability in up to 1-2 hours should be provided prior to making any commercial antimicrobial claims about surfaces with biocidal properties. US EPA guidance, originally developed for the assessment of copper surfaces, suggests antimicrobial activity assessment via application of a small volume of bacterial inoculum onto the test surface in the presence of organic soiling and subsequent dry exposure at ambient conditions [7]. Guidance documentation of the European Biocidal Product Regulation [4] suggests several testing formats at concept level but refers to only one established method, to the ISO 22196 [8]. The latter is an industrial standard that uses a low-organic liquid inoculum exposed to test surfaces as a thin layer in warm humid conditions resulting in high surface area to inoculum volume ratio. Relevance of the ISO 22196 in efficacy evaluation of surfaces developed for ambient/dry use conditions has been repeatedly questioned [9–12] and a more relevant dry method is currently under development by the ISO (ISO/DIS 7581) [13]. Experimental approaches to simulated use methods can also be found in the literature [11,14,15] but those often rely on restrictive instrumentation or methodology such as aerosolization of the inoculum. To date, there is no consensus on the combined effect of drying and exposure medium contents to antimicrobial activity of solid surfaces in conditions resembling indoor ambient environment in different microbial contamination scenarios.

In terms of efficacy assessment, several test formats have been described [9,16]. However, antimicrobial surfaces are more challenging than liquid formulations and it has been clearly demonstrated that applying different test methods can result in substantial differences in the antimicrobial activity of the surfaces. There are indications that methods involving dry exposure show lower antimicrobial activity than methods where bacterial exposure to surfaces is carried out in wet conditions. Methods also vary in the presence and amount of organic soiling as well as changes in incubation temperature during exposure. It has been demonstrated that a dry droplet and a surface transfer method results in markedly lower antibacterial activity than ISO 22196 standard test which requires exposure of bacteria to surfaces in a thin layer of liquid under a plastic film to avoid drying [17,18]. Similarly, it has been shown that antimicrobial effect of copper alloy surfaces was substantially lower in touch transfer assay than in ISO 22196 conditions [14]. One possible explanation for the lower apparent efficacy of antimicrobial surfaces in dry conditions may be the potential selection of desiccation-resistant bacteria in dry environment and co-resistance of such bacteria towards antimicrobial compounds [14]. Also, high loss of viable bacteria in negative control has been proposed [17], although quality control of antimicrobial assay should eliminate this possibility.

To date there are no systematic studies on how the different test conditions affect antimicrobial activity of solid surfaces and most of the studies have compared one or two test formats with the current standard ISO 22196 or its relevant in-house modifications. One of the widest selections of varying test protocols has been used by van de Lagemaat *et al.* who compared the antibacterial efficacy of a quaternary ammonium compound (QAC) based polymer surface using five different test methods and demonstrated method-dependent differences in antibacterial activity [18]. However, the details of the used test methods varied in many aspects, *e.g.,* in bacterial inoculum density, nutrient content and even exposure time. Therefore, the contribution of one specific test condition on antimicrobial efficacy of the tested QAC surface was very hard to reveal. Similarly, Knobloch *et al.* used in their study a specified touch transfer method where a bare finger or a glove-covered fingertip that was previously slightly moistened and contained certain level of organic soiling was used to transfer bacteria to antimicrobial surface and showed that in their assay, a series of antimicrobial surfaces were significantly less active than when tested with standard wet method ISO 22196 [14]. Although moisture and soiling of the environment appear as the most crucial factors in antimicrobial efficacy assessment, the effect of those factors was not studied in detail by Knobloch *et al*. and their contribution to the changes in antibacterial efficacy cannot be deduced from this study. Wiegand *et al.* studied a wider selection of environmental criteria affecting antibacterial efficacy but doing so only in wet test format analogous to ISO 22196 [19]. This study showed that the apparent efficacy of polyamide surfaces with zinc additives was significantly affected by the source of bacterial inoculum, inoculum density and expectedly, organic content in the exposure medium.

Due to the lack of current understanding on the role of crucial variables of test environment during dry or semi-dry antimicrobial testing of surfaces, we carried out a systematic series of antibacterial experiments in a predefined range of environmental conditions to understand, to what extent antibacterial activity of solid surfaces is determined by drying and organic soiling during exposure. We selected five solid non-porous surfaces, which had previously been included in antimicrobial studies or presented commercial antimicrobial claims. Antibacterial activity in varying test conditions was studied against two model bacteria, Gram-negative *Escherichia coli* and Gram-positive *Staphylococcus aureus.* The obtained data were expected to provide crucial information on the role of exposure conditions on antimicrobial efficacy assessment and provide insight into the possible discrepancy between claimed and application-relevant efficacy of antimicrobial surfaces.

## Materials and Methods

### Surfaces

Five test surfaces with previously shown or claimed antimicrobial effect and two control surfaces were selected: metallic copper coupons, CuC (99% Cu, Metroprint OY, Estonia); copper-based Iniesta Covidsafe adhesive tape applied to stainless steel coupons, CuT (Clean Touch Medical, Finland); metallic silver coupons, AgC (99.95% Ag, Surepure Chemetals, USA); silver-containing acrylic aqueous emulsion “TOUCH Antimicrobial coating” (silver paint) applied to stainless steel coupons, AgP (0.3-0.5 weight% silver-based organomodified bentonite, Bromoco International Ltd., UK); and a quaternary ammonium compound based Si-Quat coating applied to stainless steel coupons, SQ (active ingredient dimethyloctadecyl (3-(trimethoxysilyl)propyl) ammonium chloride (CAS 27668-52-6), Affix Labs, Finland). These test surfaces were selected to represent materials with different modes of antimicrobial action and potentially different efficiency in different humidity conditions. Two control surfaces were used: austenitic stainless steel, SS (AISI 304 SS, major elements in addition to Fe were 18% of Cr, 8% of Ni, 1.4% of Mn; 2B finish; Aperam-Stainless France;) that has been frequently suggested as a control surface in studies involving metal alloys as well as standard biocide testing methods, and borosilicate glass, NC (Corning Inc., USA) used as negative control or an inert solid surface. Additionally, polypropylene plastic, PP (Etra OY, Finland) was used as an inert cover layer on glass control in modified ISO 22196 test.

Metallic copper, silver, and steel were obtained as 1-2 mm thick sheets, that were laser-cut to 20 mm diameter round coupons. Prior to use, the metal coupons were vigorously shaken in acetone followed by a wash in deionized water and another shaking in 70% ethanol. Coupons were then air-dried in aseptic conditions and stored in sterile Petri dishes prior to use. Silver coupons were re-used during the experiment and therefore, those surfaces were always cleaned using the following procedure: a three-step shaking (2 min), sonication (15 min) and shaking (2 min) procedure in 5% citric acid, 100% acetone and 70% ethanol with 0.3 mm glass beads. Between every wash procedure, the coupons were rinsed with deionized water and after the last step, were sterilized at least for 15 min by UVC in a biosafety cabinet. A slight increase in antibacterial activity was detected after 5 cleaning cycles (Supplementary Fig S1). Based on this data, all silver coupons were subjected to 9 successive cleaning cycles before use.

CovidSafe surfaces were prepared by adhesion of ∅ 20 mm pieces of CovidSafe copper tape onto ∅ 20 mm steel coupons. Spin-coating was chosen for application of TOUCH Antimicrobial silver paint and Si-Quat coating to achieve reproducible uniform thin layer of coating on steel. For that, 100 µL of TOUCH Antimicrobial silver paint or Si-Quat coating was placed to the center of a steel coupon while spinning at 3000 rpm speed using an in-house built spin coater. Prior to use, these surfaces as well as glass and PP control surfaces were rinsed with 70% ethanol, air dried and treated for at least 15 min with UVC light in a biosafety cabinet.

### Physico-chemical characterization of surfaces

To characterize the hydrophobicity/hydrophilicity contact angles of the surfaces were measured. The sessile drop technique [20] and moving platform based on the Thorlab DT12 dovetail translation stage were used. Onto a dry surface, 2 µL drop of deionized water was pipetted in ambient conditions and after 10 seconds, water droplet was photographed using Canon EOS 650d camera equipped with an MP-E 65 mm f/2.8 1-5x macro focus lens. Using image analysis software (ImageJ 1.8.0 172 for Windows with a plugin for contact angle measurement [21]), the contact angle formed by the liquid drop on the studied surface was calculated. Four water droplets per sample, each pipetted at a different area of the surface, were used. Photos of representative water droplets for contact angle measurement are presented on Supplementary Figure S2.

X-ray photoelectron spectroscopy (XPS) as an extremely surface-sensitive technique was used to analyze the elemental composition of the topmost layer of the surfaces. All XPS measurements were conducted using Scienta SES-100 electron energy analyzer and Thermo XR3E2 non-monochromatized twin-anode X-ray source (Al K_α_ 1486.6 eV, Mg K_α_ 1253.6 eV) in ultrahigh vacuum (UHV) conditions at 10^-9^ mbar. TOUCH Antimicrobial silver paint surface was prepared for XPS by baking in the oven for 6 h at 150 °C. Al K_α_ anode with power 100 W was used for glass and TOUCH Antimicrobial surfaces, for other surfaces, power 400 W of Al K_α_ anode was applied. For all the surfaces except Covidsafe copper tape, Ar ion sputtering (5 keV, current 10 mA during 1 h) under UHV conditions at 10^-6^ mbar was performed to completely remove the surface’s topmost layer. CasaXPS software (Fairley, N. CasaXPS; Version 2.3.23; Casa Software Ltd.: Teignmouth, UK, 2000) was used for XPS data processing. Monte Carlo method [22] was used to calculate variation of XPS data.

Elemental analysis by ICP-MS was carried out with TOUCH Antimicrobial silver paint and CovidSafe copper tape. For that, a known weight of CovidSafe copper tape and a known volume of liquid TOUCH Antimicrobial silver paint formulation was dissolved in 3:1 mixture of HNO_3_: HCl following by treatment in the Berghof Speedwave Xpert microwave oven. The concentration of Ag in TOUCH Antimicrobial silver paint and Cu in CovidSafe copper tape was measured using Agilent 7700 ICP-MS from samples after their dilution up to 0.5% nitric acid.

Scanning electron microscope (SEM) (Tescan VEGA-II) with 5 kV and 20 kV electron acceleration voltages and secondary electron detector was used to visualize the top view of all the samples. Cross-sections of Covidsafe copper tape, TOUCH Antimicrobial silver paint and Si-Quat coating were prepared by cutting the coupons with respective surface coatings and removing the overhangs by a scalpel. These samples were then placed directly on the SEM stub, a narrow strip of carbon tape was placed on top of it to improve the conductivity between the surface and the SEM stub and the samples were imaged using FEI Nova NanoSEM 450 with 3 kV electron acceleration voltage. In all cases, non-conducting surfaces were covered with 10 nm layer of gold.

### Testing of antibacterial effect of the surfaces

Antibacterial effect of surfaces was analyzed after inoculation of surfaces with large droplets (50 μl) or microdroplets (5x2 μl) and exposing those droplets to different environmental variables (Table 1). For reference, surfaces were also analyzed using a modified ISO 22196 protocol.

**Table 1.**
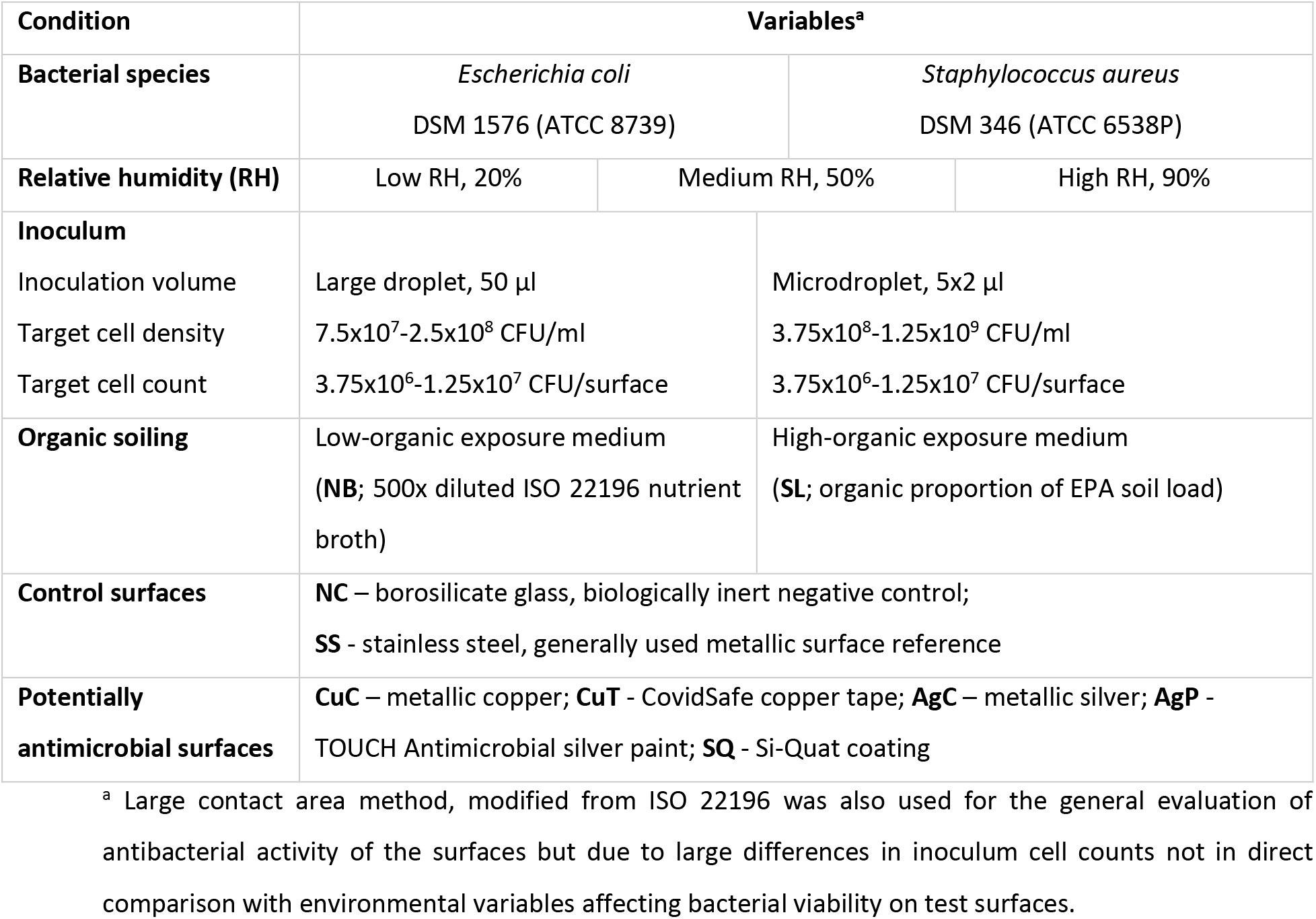
Environmental variables used in exposed large or microdroplet format antibacterial activity assessment.

Bacterial suspensions of *Escherichia coli* DSM 1576 (ATCC 8739) and *Staphylococcus aureus* DSM 346 (ATCC 6538P), both acquired from DSMZ, were prepared from fresh overnight cultures on LB solid medium (5 g/L yeast extract, 10 g/L tryptone, 5 g/L NaCl, 15 g/L agar). Bacterial biomass was suspended in sterile deionized water using a sterile inoculation loop, thoroughly mixed by pipetting and vortexing. The suspension cell counts were photometrically adjusted to target suspension densities of 1.5x10^8^-5x10^8^ or 7.5x10^8^-2.5x10^9^ CFU/ml for large droplets and microdroplets, respectively. The prepared bacterial suspension was mixed 1:1 with 2-fold concentrated organic soiling solution. The applied organic soiling was either the organic proportion of the EPA soil load (**SL**; final concentration: bovine serum albumin 2.5 g/l, yeast extract 3.5 g/l, mucin 0.8 g/l [7]) or the 500-fold diluted nutrient broth of ISO 22196 (**NB**; final concentration of ingredients: 0.006 g/l meat extract, 0.02 g/l peptone, 0.01 g/l NaCl [8] resulting in 3.75x10^6^-1.25x10^7^ CFU/surface independent of inoculum droplet size. The cell count per surface was intentionally selected higher than ISO 22196 requirement to allow assessment of at least 3 log_10_ reduction in viability in conditions that also affect viability on control surfaces and to better resemble conditions used to evaluate efficacy of surface disinfectants. Inoculum density from the European standard 13697 [23] “dirty” conditions was used as guidance for large droplets and the same cell count per surface was also used for microdroplet inoculation with higher inoculum density. [7]

Surfaces with large 50 μl droplets or 5x2 μl microdroplets on surfaces were incubated in open Petri dishes (Supplementary Fig S2) in a climate chamber (Climacell EVO, Memmert, USA). Incubation was carried out at 22°C and variable RH: 90% RH (high humidity), 50% RH (moderate humidity) or 20% RH (low humidity). After 0.5, 1, 2 and 6 h of incubation the surfaces submerged to 10 mL of toxicity neutralizing medium (SCDLP: 17 g/L casein peptone, 3 g/L soybean peptone, 5 g/L NaCl, 2.5 g/L Na_2_HPO_4_, 2.5 g/L glucose, 1.0 g/L lecithin, 7.0 g/L Tween80 [8]) in 50 mL conical centrifuge tubes. The tubes were vortexed for 30 seconds to detach bacteria, serially diluted in phosphate-buffered saline (10-fold stock diluted to: 8 g/L NaCl, 0.2 g/L KCl, 1.44 g/L Na_2_HPO_4_, 0.2 g/L KH_2_PO_4_; pH 7.1). Dilutions were drop-plated, incubated at 37°C for optimal growth and plate counts registered after 16-18 h for *E. coli* or 24 h for *S. aureus.* At least 3 biological and 2 technical repeats were analyzed.

For reference and comparison, all the surfaces were tested using a modification of ISO 22196 format. For that, inoculums of *E. coli* and *S. aureus* were photometrically adjusted to the target 3.14x10^6^ CFU/ml in 500-fold diluted NB broth and 15 µL of this inoculum (ca 1.5x10^4^ CFU/cm^2^; 4.7x10^4^ CFU/surface) was pipetted onto each surface. The surfaces were then covered by 25x25 mm coverslips with an overhang but avoiding inoculum leakage and incubated at room temperature (22°C) in a >90% RH environment for 0.5, 1, 2 and 6 h. After exposure, the surfaces were placed into 10 ml of SCDLP neutralizing medium, cells were recovered and plated for CFU counts as described above. These experiments were performed in three biological replicates.

### Statistical analysis

Statistical analysis of the data was performed with GraphPad Prism 9.5.0 (GraphPad Software, San Diego, USA). Raw data used can be found in the Supplementary Raw Data file. Data from the 2 h time point was selected for a more thorough statistical analysis as longest relevant contact time in the context of current legislative framework in the US and EU. Log_10_-transformed data was used for the analysis and values below limit of quantification (LOQ) were brought to LOQ prior to analysis. LOQ was considered as at least 3 colonies counted from of 20 μl or 500 μl undiluted surface wash-off plated from droplet inoculation or ISO format, respectively, resulting in LOQ of 3.18 or 1.48 log_10_CFU/surface. One-way, two-way, and three-way ANOVA followed by Tuckey or Dunnett *post-hoc* tests at α=0.05 were used where appropriate.

## Results and discussion

### Characteristics of the surfaces

The physical appearance of the surfaces studied is shown in Figures 1 and 2. Metallic copper and silver were relatively smooth at their pristine state. Differently from copper and silver, other surfaces demonstrated certain surface structures already in their pristine form. CovidSafe copper tape exhibited a clear surface structure (Fig 1, CuT) and from cross-section (Fig 2, CuT) its thickness was estimated to be 5-6 µm. Both TOUCH Antimicrobial silver paint and Si-Quat coating were coated onto stainless steel coupons. The steel itself exhibited clear surface structures (Fig 1, SS) and as those structures were well visible in case of Si-Quat coating (Fig 1, SQ), we concluded that Si-Quat coating was relatively thin allowing secondary electrons from the stainless steel pass through the coating layer. This was confirmed also from the cross-section of Si-Quat coated samples, where the thickness of the coating was calculated to be 3-5 μm. Unlike Si-Quat, no steel surface structures were visible with SEM in case of TOUCH Antimicrobial silver paint (Fig 1, AgP). According to cross-section (Fig 2, AgP), the thickness of TOUCH Antimicrobial silver paint on steel was reaching 10 µm and was therefore able to mask the surface structure of steel. Glass surface appeared completely smooth as expected (Fig 1, NC).

**Figure 1.**
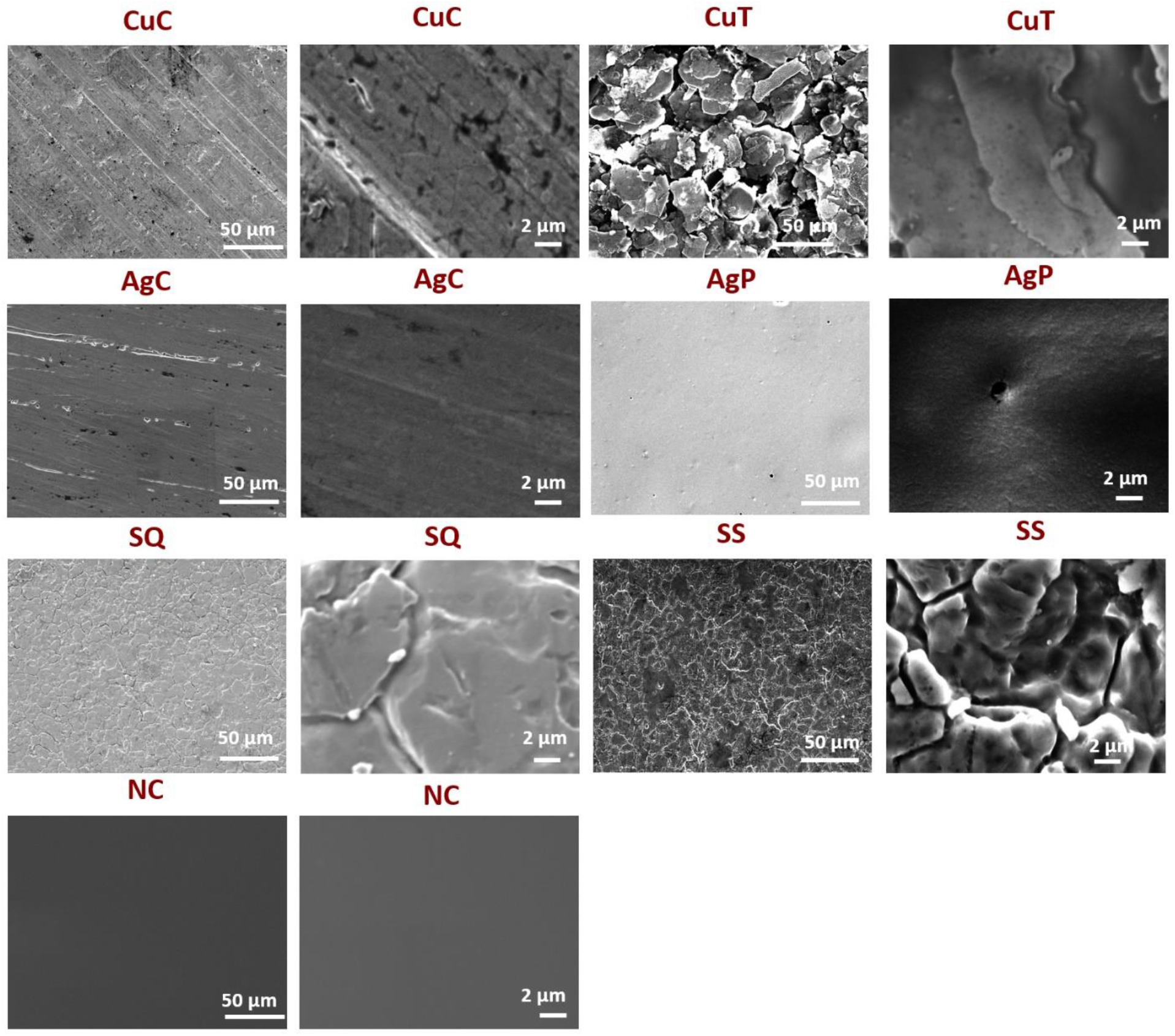
SEM images of the studied surfaces. (CuC – metallic copper; CuT – CovidSafe copper tape; AgC – metallic silver; AgP – TOUCH Antimicrobial silver paint; SQ – Si-Quat coating, SS – stainless steel; NC – borosilicate glass) in small (left) and large magnification (right).

**Figure 2.**
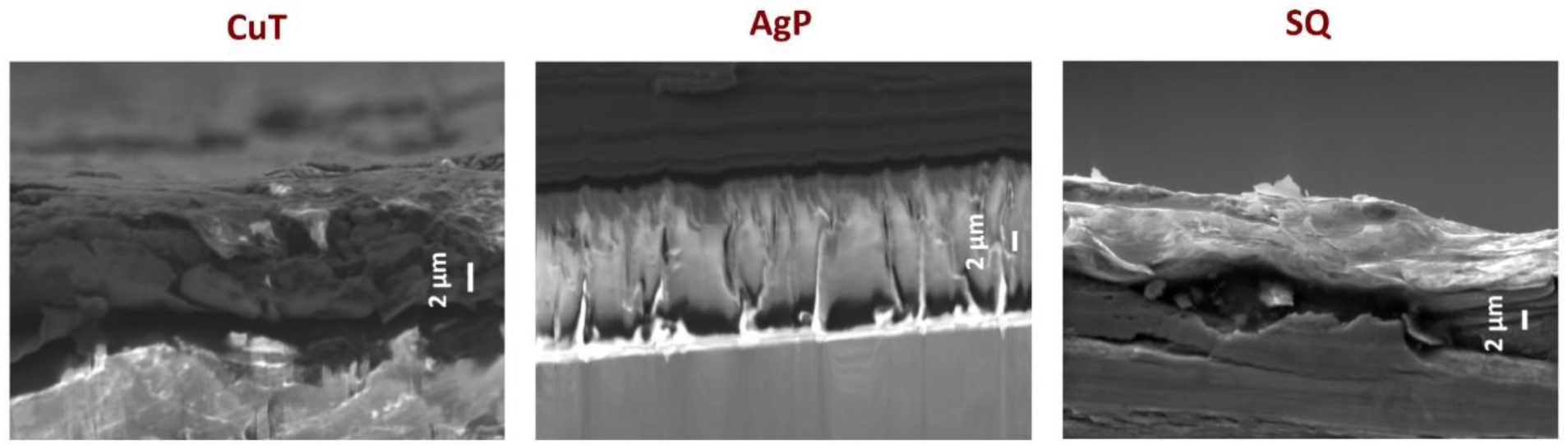
Cross sections of CuT (CovidSafe copper tape), AgP (TOUCH Antimicrobial silver paint) and SQ (Si-Quat coating) on stainless steel under SEM. In case of CovidSafe, both, copper-containing (darker) surface layer and adhesive polymer (lighter) base layer can be seen.

According to XPS, metallic copper and silver presented the respective elements Cu and Ag on their very surface (Table 2) as expected. However, since XPS is an extremely surface sensitive method (surface depth up to 10 nm) significant amount of O and C were present in XPS spectra as surface contaminants. Such a situation is typical for XPS measurements. To reveal the composition of copper and silver inside samples, Ar^+^ ion sputtering was used to remove uppermost sample layer prior to XPS analysis, which revealed that metallic copper and silver were presenting 100% Cu and 93% Ag on their surfaces respectively (Table 2). On stainless steel, almost all expected metallic components (Fe, Cr and Mn) were identified. Nickel was not detected with XPS but given that this is a surface sensitive method, we suggest that it (8% Ni content in AISI 304 stainless steel) may be not present on the very first surface layers of the 2B finish.

**Table 2.** Characteristics of the surfaces used in the study.

| Surface | Composition |  |  |
| --- | --- | --- | --- |
|  | ICP-MS | XPS | XPS after Ar<sup>+</sup> sputtering |
| **CuC**; 99% metallic **copper**<br>(Metroprint OY, Estonia) | n.a. | 56±0.41% C, 25±0.28% O,<br>**14±0.17% Cu**, 5±0.16% N | **100±0.55 % Cu** |
| **CuT**; Iniesta **Covidsafe** copper tape<br>(Clean Touch Medical, Finland) on<br>stainless steel | 100 000 mg Cu<br>per kg of tape | n.a. | n.a. |
| **AgC**; 99.95% metallic **silver**<br>(Surepure Chemetals, USA) | n.a. | 58±0.41% C , **25±0.74%**<br>**Ag**, 18±0.72% O, | **93±0.69 % Ag**, 3±0.48% O,<br>4±0.54% C |
| **AgP**; **TOUCH Antimicrobial** silver<br>paint (TOUCH Antimicrobial, UK) on<br>stainless steel | 27 mg Ag in kg<br>of paint | 73±0.57% C, 22±0.55% O,<br>6±0.26% Si, < **1% Ag** | 98±0.41% C, 2±0.39% O, <<br>**1% Ag** |
| **SQ**; **Si-Quat** coating (Affix Labs,<br>Finland) on stainless steel | n.a. | 67±0.37% C, 20±0.22% O,<br>10±0.33% Si, 2±0.15% Cl,<br>1±0.20% N | 70±1.53% Fe, 14±0.37%<br>Cr, 6±0.93% C, 5±0.24% O,<br>5±0.17% Mn |
| **SS**; austenitic stainless **steel**<br>coupons; AISI 304 SS; finish 2B<br>(Aperam- Stainless, France) | n.a. | 47±0.53% O, 40±0.51% C,<br>**7±0.42% Fe**, **4±0.20% Cr**,<br>**1±0.14% Mn**, < **0.5% Ni** | **67±0.57% Fe**, **12±0.18%**<br>**Cr**, 9±0.25% O, 6±0.57% C,<br>**6±0.11% Mn**, < **0.5% Ni** |
| **NC**; borosilicate **glass** (Corning Inc.,<br>USA) | n.a. | 67±1.11% C, 25±0.92% O,<br>8±0.84% Si | n.a. |

n.a. not analysed

Although Ag was expected to be present on TOUCH Antimicrobial silver paint surface, XPS was not able to reveal any elemental Ag on the surface layer of that sample (Table 2). To eliminate the possibility that Ag was masked by surface contaminants such as C or O, TOUCH Antimicrobial surface was cleaned using Ar^+^ ion sputtering. Interestingly, no Ag was detected also on ion-sputtered samples (Table 2) suggesting that the concentration of Ag was below the limit of determination of the method (< 1% Ag). To prove the presence of Ag in the sample, ICP-MS analysis of the original liquid TOUCH Antimicrobial coating formulation was performed and 20 mg of Ag /kg of the formulation was identified (Table 2).

On Si-Quat surface, Si, Cl and N were detected, which corresponds to the chemical formula of the active compound in Si-Quat coating (C_26_H_58_NO_3_ClSi) (Table 2). Ar^+^ ion sputtering of Si-Quat surface removed most of the Si-Quat coating as in ion-sputtered samples no Si or Cl was detected and instead, components of the base material, stainless steel (Fe, Cr and Mn) were revealed (Table 2).

Due to the outgassing from the adhesive, UHV conditions in the analysis chamber were not achievable and CovidSafe copper tape could not be safely analyzed with XPS without the risk of damage to the instrument. To prove the presence of Cu on that surface, ICP-MS analysis was performed on the bulk CovidSafe material, which proved the presence of Cu in that material (Table 2).

Most of the surfaces were hydrophilic in nature with just two surfaces – CovidSafe and steel having slightly hydrophobic properties with contact angles >90° (Fig 3). Similar behavior of bacterial inoculum droplets was observed on all surfaces (Supplementary Fig S2) and thus, in antibacterial tests, differences in hydrophilicity of surfaces was not considered important. However, uneven drying time of 2 μl microdroplets on the same surface was often observed, possibly explained not solely or mainly by the physical characteristics of the surfaces but also ventilation in the climate chamber, variation in pipetting or combination of the above.

**Figure 3.**
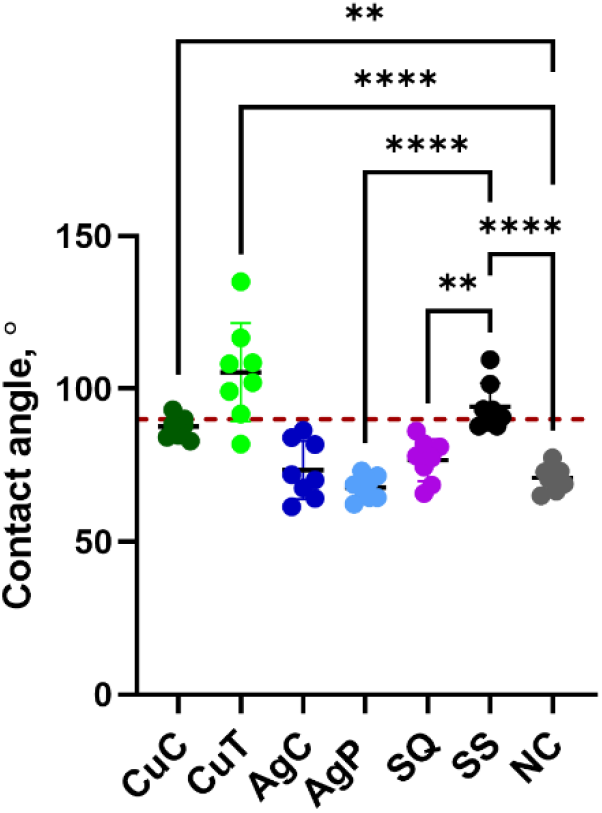
Contact angle measurements of the studied surfaces. (CuC, metallic coupon; CuT, CovidSafe copper tape; AgC, metallic silver; AgP, TOUCH Antimicrobial silver paint; SQ, Si-Quat coating, SS, stainless steel, NC, borosilicate glass). The coupons were rinsed with acetone and ethanol the same way as prior to antibacterial testing. Statistically significant differences (P<0.05) from the NC and SS control surfaces are marked with **** (P≤ 0.0001), ** (P≤ 0.01). The 90° boundary above which the values can be considered hydrophobic, is marked with a red dotted line.

### Effect of test format on antibacterial activity

Antibacterial activity of the five potentially antimicrobial surfaces compared to 2 control surfaces was evaluated in two test formats: an environmentally exposed large droplet or microdroplet inoculation in various semi-dry conditions and a modified ISO 22196 [8] wet test as an industrial reference. Exposure in semi-dry conditions was selected to resemble mucin-containing oral/nasal spray and spray-contamination of surfaces or larger droplet contamination with low organic content e.g., near faucets or cooking surfaces, on surfaces in public space, non-invasive surfaces in health-care settings or in the food industry. As a widely used industrial reference, ISO 22196 format employing large surface area to inoculum volume ratio was used in otherwise similar conditions to test the maximum expected efficacy of the surfaces. 500-fold diluted nutrient broth (0.026 g/L organic content) with relatively low metal ion complexing properties was chosen as a low-organic exposure medium also used in ISO 22196. The organic proportion of mucin-containing soil load described in US EPA test guidance, originally developed for the efficacy assessment of copper surfaces [7] was chosen as a high-organic exposure medium (6.8 g/L organic content) that has metal ion complexing capacity and supports metabolism. PBS was omitted from the EPA SL formulation to avoid introducing additional osmotic variable into the comparison of environmental conditions. In addition to organic soiling, relative air humidity (RH) was varied between exposures, to represent dry (20%), normal (50%) or humid (90%) indoor conditions. The test surfaces were generally chosen due to their proven antibacterial efficacy. Copper and silver were included as materials with historically well-known antimicrobial properties as well as continuing research interest and commercial use [24]. Quaternary ammonium-based Si-Quat coating was selected as a non-metallic surface with a different mode of action. Antibacterial activity of those surfaces was evaluated in comparison with borosilicate glass as inert surface and stainless steel as a widely used non-antimicrobial metal surface.

Figures 4 and 5 present the log_10_ reduction in viability of *E. coli* and *S. aureus*, respectively, on studied surfaces after 2 h exposure in all the test conditions used (complete data from 0.5-6 h time points: Supplementary Fig S3, S4 and Supplementary Tables S1, S2). Copper-based surfaces proved to be the most efficient antibacterial surfaces across the conditions tested, followed by silver and Si-Quat surfaces (Fig 4, 5) with the latter showing potential specifically towards *S. aureus* (Fig 5). The CovidSafe copper tape demonstrated lower effect than pure copper and the TOUCH Antimicrobial silver paint was the least effective with no effect in any of the test conditions in 2 h (Fig 4, 5) and only a minor effect against *E. coli* in a couple of conditions by the longest 6 h time point.

**Figure 4.**
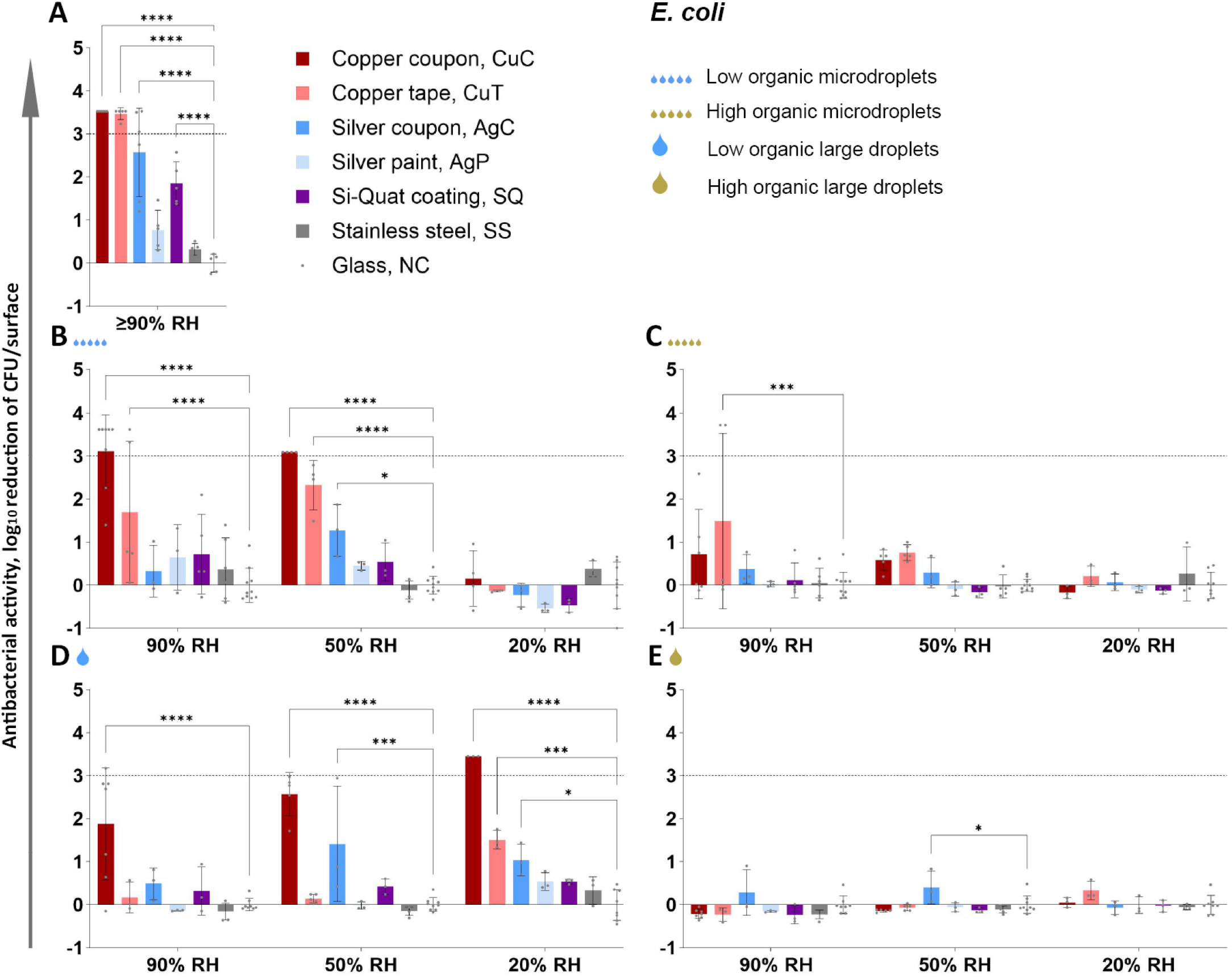
Antibacterial activity of the surfaces towards *E. coli* after 2 h exposure in low organic (A, B, D) and in high organic (C, E) media exposed either as a liquid layer ISO 22196 format (A), microdroplets (5x2 μl; B, C) or large droplet (50 μl; D, E). Log_10_ reduction is calculated from glass (NC). The maximum possible log_10_ reduction detectable depended on bacterial viability on glass control surface in each test condition. Maximum log_10_ reduction was achieved by copper coupon (CuC) on panel A, B (50% RH) and D (20% RH) indicated by high antibacterial activity values with no error bars. An average of 3-6 biological replicates with standard deviation is shown. Statistically significant difference (P<0.05) from the NC control surface is marked with **** (P≤ 0.0001), *** (P≤0,001), ** (P≤ 0.01), * (P<0.05). The target of 3 log_10_ reduction is displayed as a grey dotted line. Antibacterial activity after 0.5, 1, 2 and 6 h exposure is presented on Supplementary Figures S3 and S4, log_10_-transformed viable counts in Supplementary Tables S1 and S2 and raw data in Supplementary Raw Data file.

**Figure 5.**
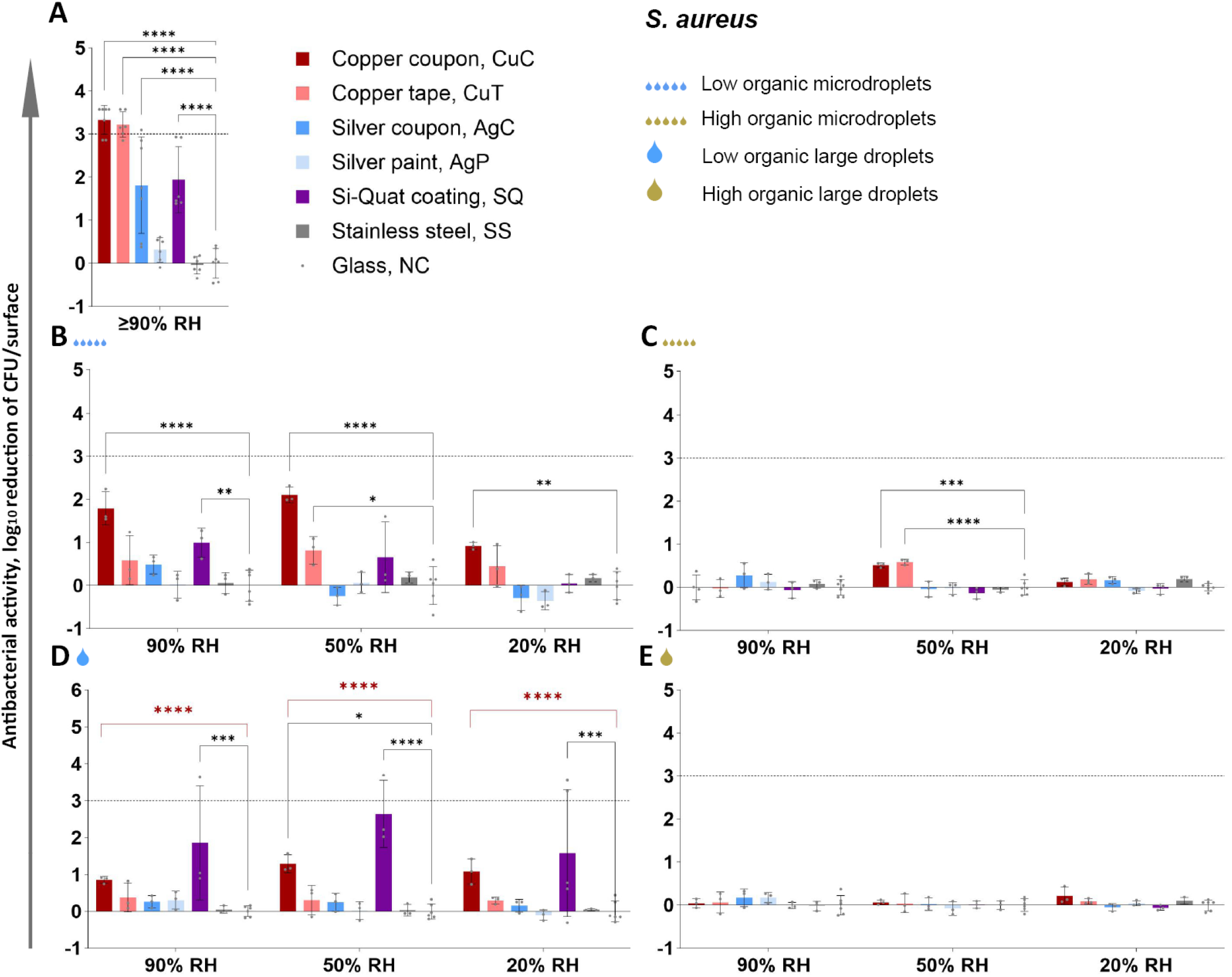
Antibacterial activity of the surfaces towards *S. aureus* after 2 h exposure in low organic (A, B, D) and in high organic (C, E) media exposed either as a liquid layer in ISO 22196 format (A), microdroplets (5x2 μl; B, C) or large droplet (50 μl; D, E). Log_10_ reduction is calculated from glass (NC). The maximum possible log_10_ reduction detectable depended on bacterial viability on glass control surface in each test condition. Maximum possible log_10_ reduction of *S. aureus* was not achieved by any of the surfaces in any conditions after 2 h exposure. An average of 3-6 biological replicates with standard deviation is shown. Statistically significant difference (P<0.05) from the NC control surface is marked with **** (P≤ 0.0001), *** (P≤0,001), ** (P≤ 0.01), * (P<0.05). On panel D, two-way ANOVA + post-hoc results are shown for the whole dataset (black *-***) as well as with the most variable SQ group removed (red ****). The target of 3 log_10_ reduction is displayed as a grey dotted line. Antibacterial activity after 0.5, 1, 2 and 6 h exposure is presented on Supplementary Figures S3 and S4, log_10_-transformed viable counts in Supplementary Tables S1 and S2 and raw data in Supplementary Raw Data file.

Large contact area between the bacterial inoculum and the test surface in humid conditions in the modified ISO 22196 method (Fig 4A, 5A) generally overestimated antibacterial activity of metal-based surfaces compared to droplet and microdroplet inoculation in the same low-organic exposure medium and at ambient temperature (panels B and D on Fig 4 and 5). The precise log_10_ reduction values are challenging to compare due to substantially lower cell count per surface in the ISO format as that can affect the antibacterial effect size. The ISO format also expectedly guaranteed sufficient bacterial viability on control surfaces (Supplementary Tables S1, S2; Fig 7) which simplifies analysis in a standard test format but does not reflect application-relevant conditions where a substantial amount of bacteria might be killed or inactivated by environmental variables *e.g*., desiccation, light exposure, organic and non-organic surface contamination, too much or too little oxygen etc. It could be that combinations of environmental variables might decrease bacterial viability synergistically and complicate efficacy assessment of antimicrobial surfaces. For example, we have previously reported that low-intensity UVA phototoxicity towards both *E. coli* and *S. aureus* is higher in lower air humidity on inert control surfaces [25]. Although we did not specifically design the current study to quantify synergism between the selected environmental parameters, there is clear evidence of synergistic effects represented by substantial contribution of statistically significant 2-way and 3-way interactions of environmental parameters to total variability of viable counts based on 3-way ANOVA of 2 h exposure data (Fig 6).

**Figure 6.**
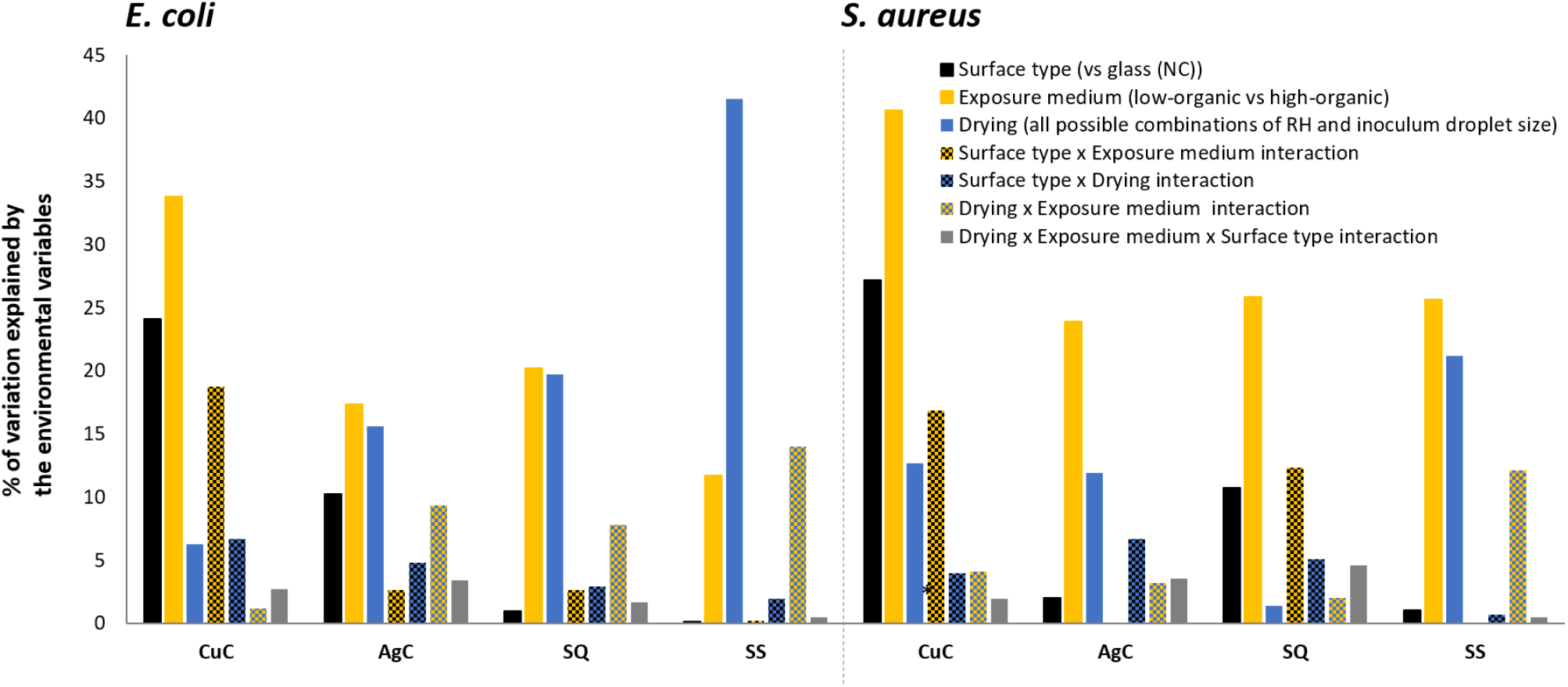
Contribution of drying of the inoculum, surface type (*vs* glass control, NC), exposure medium (high vs low organic content) and their interactions (indicated by variable x variable interaction in the legend) to the total variation in viable counts. based on 3-way ANOVA analysis of 2 h exposure data of *E. coli* (left) and *S. aureus* (right). The parameters of relative humidity (RH) and inoculum droplet size proved to be interdependent in their effect on viability probably by affecting drying time and could thereby impair main effect and interaction interpretation in multifactorial ANOVA. Instead, a combined variable, drying, was used. As a categorical variable drying comprised of all 6 possible combinations of RH (20%, 50% and 90%) and inoculum droplet sizes (5x2 μl and 50 μl). Only the most effective surface type of each active agent and stainless steel as a metallic control surface are shown compared to glass (NC): CuC, metallic copper; AgC, metallic silver coupon; SQ, Si-Quat coating; SS, stainless steel. All values above 6% are statistically significant. Numerical data is presented in Supplementary Table S3.

**Figure 7.**
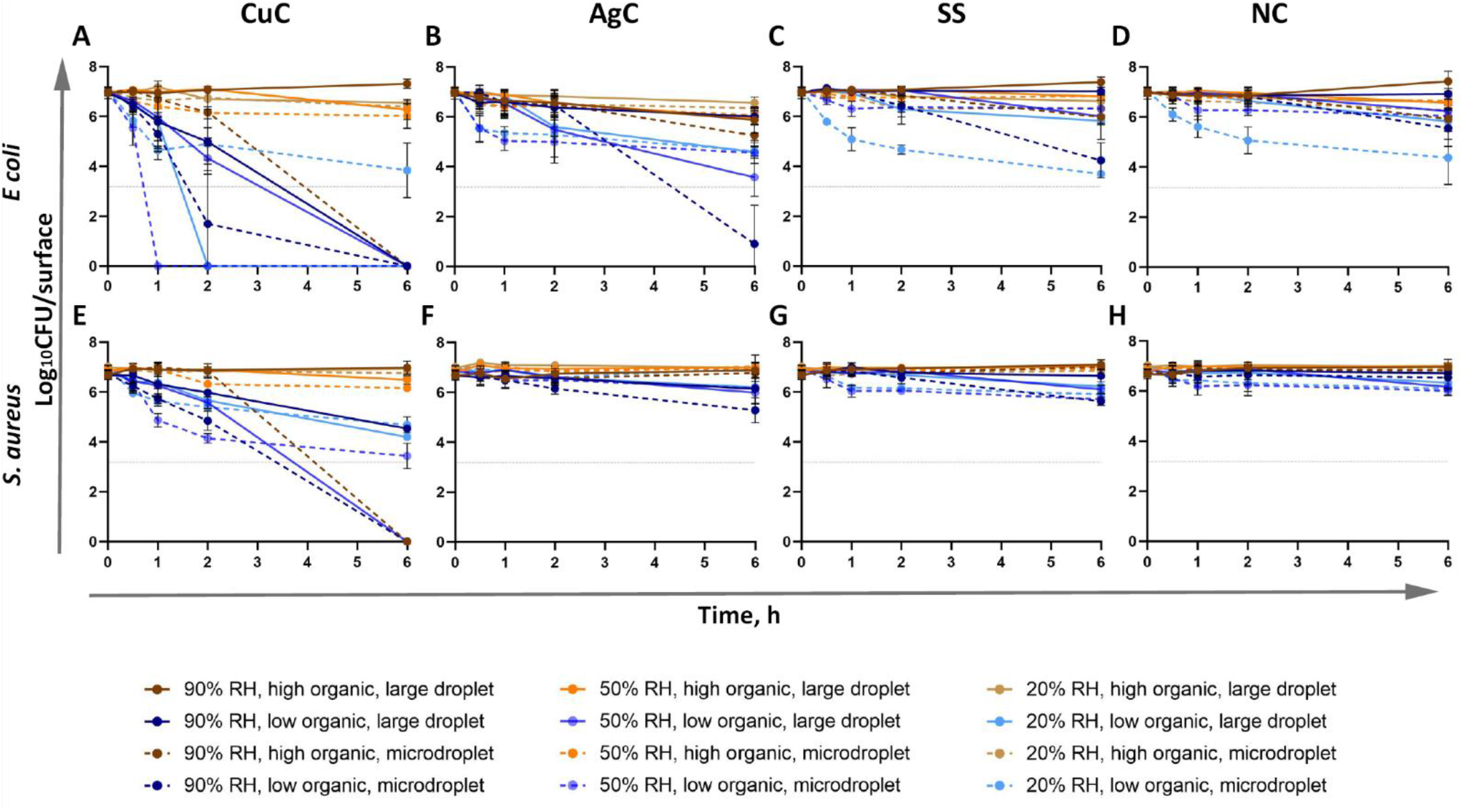
Comparison of antibacterial effect of the most effective surface types and control surfaces towards *E. coli* (A-D) and *S. aureus* (E-H) in variable testing conditions. CuC, metallic copper; AgC, metallic silver; SS, stainless steel; NC borosilicate glass. Blue hues denote low and yellow hues high organic content in exposure media. Lighter color indicates lower air humidity and dotted lines microdroplet inoculation. An average of 3-6 biological replicates with standard deviation is shown. Values at or above the detection limit of at least three colonies counted in all undiluted plated drops (dotted line at y=3.18) were used for the statistical analysis of CFU counts. Lower values are shown to demonstrate that growth of viable bacteria was detected in at least some repetitions. Tabular presentation of mean values and standard deviations is presented in Supplementary Tables S1 and S2 and raw data in Supplementary Raw Data file.

Therefore, the ISO format represents the “best-case scenario” with high bacterial viability on control surface and high antimicrobial activity on surfaces of interest. With this study we demonstrate that in more plausible “worst-case scenarios” viability of *E. coli* is decreased on control surfaces and antibacterial activity of the potential antimicrobial surfaces is generally lower than in the ISO format. However, as can be seen for *E. coli* in 50% and 90% RH and microdroplets (Fig 4B) or large droplet in 20% RH (Fig 4D) in low-organic medium there can be exceptions, where antibacterial activity of copper after 2 h exposure does not significantly differ between the ISO format (-3.5 log_10_) and droplet inoculations (-3.1, 3.1 and 3.45 log_10_, respectively; P>0.05).

Although several surfaces elicited >3 log_10_ reduction in bacterial viability in different test conditions after 6 h exposure, the regulatory goal of at least 3 log_10_ reduction in up to (1-)2 h was in most cases not met, except copper-based surfaces in some conditions specified below. Copper and silver surfaces were generally more effective against *E. coli* than *S. aureus* (Fig 4, 5) while Si-Quat surfaces tended to preferentially act better on *S. aureus* (Fig 5B, 5D). However, despite substantial mean log_10_ reduction Si-Quat surfaces did not demonstrate a reliable effect due to extremely high variability among biological replicates. It must be noted that while the CovidSafe copper tape elicited >3 log_10_ reduction in viability towards both species after 2 h in the ISO format (Fig 4A, 5A), it had considerably lower to no activity in other formats, *e.g.* <1 log_10_ reduction of *S. aureus* regardless of exposed droplet test conditions. The TOUCH Antimicrobial silver paint did not show any antibacterial activity in any condition towards either of the species after 2 h exposure including the ISO 22196 format. This lack of effect could be explained by the extremely low silver content in the TOUCH Antimicrobial initial formulation and no Ag detected by XPS on the coated surfaces (Table 2). After 2 h exposure, only copper-based surfaces reached the required 3 log_10_ reduction in viability. Both metallic copper and CovidSafe copper tape demonstrated >3 log_10_ reduction of viable counts of both bacterial species in the ISO format (P<0.0001 in all cases). However, only metallic copper caused >3 log_10_ reduction in exposed droplet format only towards *E. coli* and only in the following conditions: low-organic microdroplets in 50% and 90% RH and low-organic large droplets in 20% RH (P<0.0001 in all cases).

### Effect of environmental variables on antibacterial activity

All the antimicrobial surfaces were generally less effective in dry conditions (20% RH) and/or in high-organic medium. No biologically (> 1 log_10_ reduction in viable counts compared to controls) and/or statistically significant antibacterial activity of any of the surfaces, in any of the conditions towards either of the bacterial species was observed after 2 h exposure in case of microdroplet inoculation and low relative humidity (panels B-C on Fig 4 and 5). There were also no surfaces that could biologically and statistically significantly reduce bacterial viability after 2 h in large droplets of exposure medium with high organic content (Fig 4E, 5E). Both findings highlight an urgent need to use application-relevant methodology to adequately assess efficacy of antimicrobial surfaces used in environments exposed to air.

The environmental variables affected viability of both *E. coli* and *S. aureus* on control surfaces (Fig 7C, 7D). As a general rule, on surfaces with lower to no known antibacterial activity, viability of *E. coli* was more affected by drying analyzed as a combined effect of RH and inoculum droplet size (Fig 6, left) while *S. aureus* was more affected by the selection of exposure medium (Fig 6, right). Our results indicated that the effect of environmental variables depended also on the surface type (Fig 6; based on 3-way ANOVA of 2 h exposure data). While metallic copper surfaces demonstrated the highest antibacterial activity, we can see that its effect was most dependent on exposure medium, surface type and their interaction jointly explaining 77% and 85% of total variation in viable counts for *E. coli* and *S. aureus* on copper, respectively (Fig 6).

### Effect of drying on antibacterial activity

Drying had only a minor contribution to overall variability in viable counts on copper (Fig 6), thus not dramatically affecting bacterial viability. However, drying, exposure medium and their interaction had a more even contribution to variability of viable counts on silver surface (Fig 6). Based on literature, silver surfaces are expected to have a larger effect in wet conditions and copper also demonstrating substantial effect in dry conditions [14,26,27]. This was with some reservation confirmed for *E. coli*, but not for *S. aureus,* partly because silver had overall very low antibacterial activity towards *S. aureus*. Interestingly, while metal-based surfaces seemed more efficient towards *E. coli*, Si-Quat coating preferentially worked against *S. aureus* in low-organic medium and more importantly, its activity was not significantly affected by RH as none of the drying-related causes statistically significantly contributed to viable count variation (Fig 6, right). Viable counts retrieved from Si-Quat coating in exposed droplet format were also characterized by extremely high variability compared to other surfaces, possibly indicating the unstable nature of microbe-surface interaction parameters beyond our selected criteria. The Si-Quat coating is described as a contact-killing and not release-based surface by the producer. Based on our results it seems that Si-Quat surfaces could be most effective in low-organic large droplets independent of RH conditions (Fig 5D). This in combination with our finding on higher antibacterial activity of Si-Quat coating towards *S. aureus* than towards *E. coli* suggests the possible crucial role of water environment and potentially electrostatic interactions with non-motile *S. aureus* as opposed to motile *E. coli*.

Interestingly, in the ISO 22196 format, the antimicrobial effect, if present, generally reached its maximum by 2 h and plateaued thereafter whereas substantial decrease in bacterial viability between 2 and 6 h could be observed on several surfaces in case of environmentally exposed droplets (Fig 7A, 7B; Supplementary Fig S3 and S4). In case of exposed droplets, such plateauing of the antibacterial effect was seen after complete drying of inoculum droplets (Table 3, Supplementary Fig S3 and S4). There seems to be a balance between the effect size of antibacterial action and the duration of the physical process of drying which can cause higher antibacterial activity in lower RH if the droplets do not dry so fast as microdroplets in 20% RH, possibly due to longer wet contact time, longer active agent release and eventually drying i.e., decreasing droplet volume while increasing active agent concentration (*e.g.* Fig 7A, 7B ; blue lines, both drop sizes). Plateauing of the antibacterial effect after drying could indicate that at least some amount of free water is needed to elicit the antibacterial activity. In this light the normal indoor RH (50%) provides conditions that, at least in low-organic environment, could benefit most of the use antimicrobial surfaces. However, the process of drying is not as simple as just the evaporation of water and should be studied further in the context of antimicrobial surfaces. Not only RH but also the physical characteristics of surfaces, microbes, and liquids can impact the kinetics of drying and may potentially alter the antibacterial properties of surfaces. [28–32].

**Table 3.** Drying time of the inoculum droplets. Time point (h) by which the inoculum droplets had visibly completely dried on the studied surface, irrespective of organic soiling or bacterial species used. In the case of 5x2 μl microdroplet inoculation where uneven drying time of individual droplets on the same surface, the time-point after which all droplets had visibly dried was registered. Representative images of the droplets are presented in Figure S5.

|  | 20% RH | 50% RH | 90% RH |
| --- | --- | --- | --- |
| Large droplet (50 μl) | 6 h | 6 h | >6 h |
| Microdroplets (5x2 μl) | 0.5 h | 0.5-1 h | 6 h |

### Effect of exposure media on antibacterial activity

Organic soiling is generally expected to reduce antimicrobial activity of surfaces. In this study organic proportion of soil load formulation from the EPA method was as used. The individual components of soil load are bovine serum albumin (BSA), yeast extract and mucin, which have been individually used for various purposes. BSA or fetal bovine serum (FBS) have often been used to mimic “dirty” conditions in various standard protocols and literature sources (e.g., EN 13697 [23]) but published results are contradictory. For example, BSA has been demonstrated to protect bacteria against light-activated antimicrobial surfaces [33] but also increase efficacy of copper surfaces exposed to BSA-containing bacterial aerosols [11]. Yeast extract has been shown to decrease the toxicity of heavy metals such as cadmium [34]. The heavily glycosylated mucins in respiratory droplets may offer some protection against inactivation by drying in case of enveloped viruses [35] that could be considered somewhat similar to Gram-negative bacteria with their environmentally exposed outer membrane.

The behavior of one and the same surface in different conditions is illustrated in Figure 7. Lines for high-organic conditions (brown/yellow) are generally grouping together to the top of the graphs irrespective of RH clearly indicating that antibacterial activity of the surfaces is more affected by the choice of exposure medium than drying of the inoculum. None of the test surfaces elicited any substantial antibacterial activity towards either bacterial species in high-organic exposure medium in low to medium RH. However, these are the conditions often encountered in real applications of commercial antimicrobial surfaces when we think of surfaces contaminated by bodily fluids or the food industry for example. High organic content also demonstrated a protective effect against desiccation effects in microdroplet inoculation, especially in low RH conditions where otherwise a substantial decrease in viable counts was also observed on control surfaces (Fig 7C, 7D).

Although higher efficacy than presented on Figure 7 (brown/yellow on panels A,E) definitely can be achieved in high-organic EPA soil load for certain copper-based surfaces as such products have been approved for commercial use in the US employing the EPA method [7] to back the antimicrobial claims, it has to be noted that the method instructs to smear the inoculum over a large area on the test surface. We used non-smeared droplets to intentionally mimic contamination via spray or semi-wet cross-contamination. Earlier studies have shown that copper-based surfaces elicit lower antibacterial activity when surfaces are inoculated as undisturbed microdroplets as opposed to smearing the inoculum [37]. In our previous studies, we have seen similar effect and noted that bacteria survived better in 50 μl or 75 μl droplets of LB on copper surface than in 25 μl [38]. The latter also resulted in higher copper release from the surface during the 1 h exposure. Indeed, also in the current study, copper surfaces tended to be most efficient in case of microdroplet inoculation and high RH (Fig 4B, 5B), especially in the longest time-point and highest RH where the microdroplets had not completely dried on the surfaces (Supplementary Fig S3, S4). Due to the test format and long exposures in dry conditions, we could unfortunately not determine the amount of released copper. It is also possible that differences in viability can at least partly arise from different flow dynamics and surface tension in the droplets of different exposure media during drying [32].

As the content of the EPA soil load [7] is complex, we further aimed to clarify whether it is the total organic content or one of the individual organic components of the soil load medium that protects bacteria from the antibacterial as well as from drying effects. For that, we studied the antibacterial effect of the most effective metallic surface (copper) in comparison with stainless steel in the presence and absence of soil load or its components.

Figure 8 reveals that in moderate drying conditions (50% RH) and microdroplet inoculation in case of which microdroplets dry by the end of 1-hour exposure (Table 3) viability of *E. coli* on copper was clearly dependent on organic content of the exposure medium. No decrease in viability on copper surface (P>0.05) was detected in the presence of soil load representing high-organic environment and its 10-fold dilution (Fig 8). The same was true when the components of soil load were used separately, indicating that any of the organic components in soil load is more than sufficient in eliminating the antibacterial activity of copper surfaces in moderately dry conditions. Only 100-fold diluted soil load enabled copper to cause significant reduction in viable count (-3.6 log_10_; P<0.0001) that was relatively similar to that induced by a low-organic exposure medium (-4.8 log_10_; P<0.0001). While high organic content generally reduces antimicrobial activity of metal-based agents, exceptions can be found where the opposite is claimed. Contradicting our results with 3g/L BSA in microdroplets, copper-based surfaces have also been shown to present faster antibacterial activity when exposed to aerosols containing microbes and 2.5 g/L BSA [11]. To put the organic content in the EPA soil load (6.8 g/L) into a context, the commonly used lysogeny broth (LB; [36]) often used in laboratory conditions as a growth-supporting exposure medium contains double the amount, i.e., 15 g/L of organic ingredients.

**Figure 8.**
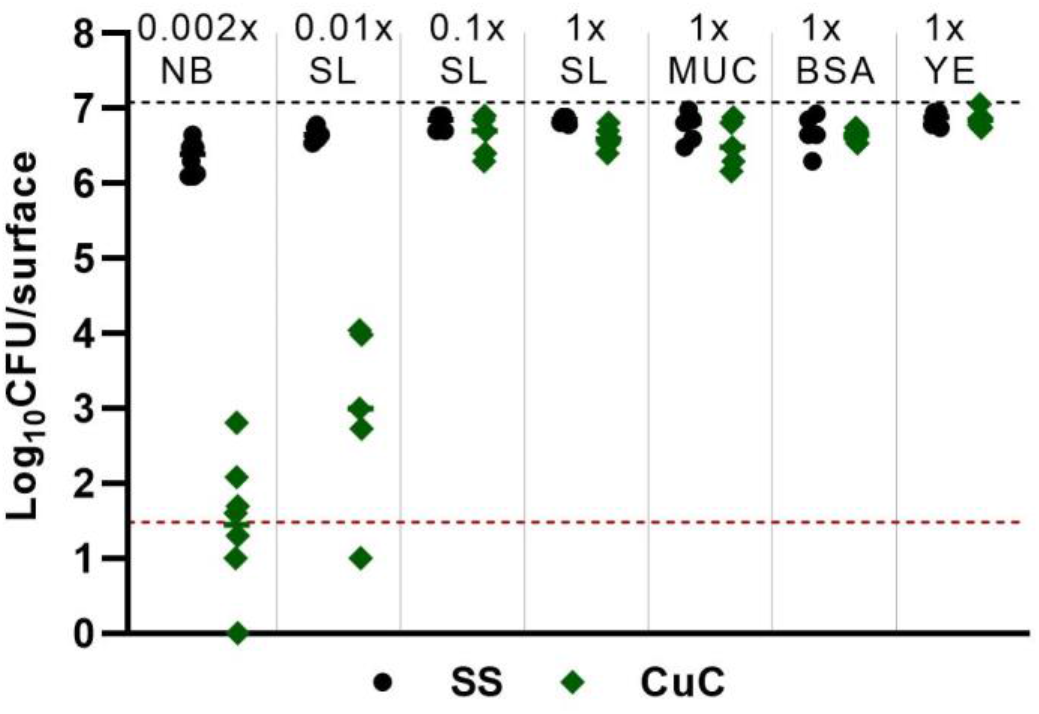
Effect of exposure medium, its organic content and individual organic soiling components on antibacterial activity of copper surfaces (**CuC**, green) against *E. coli* after 1 h exposure in microdroplets at 50% relative humidity compared to stainless steel (**SS**, black). Organic soiling scenarios tested: 500-fold diluted nutrient broth from ISO 22196 (**0.002x NB**; 0.026 g/L organics), 100-fold diluted EPA soil load (**0.01x SL**; 0.068 g/L organics), 10-fold diluted EPA soil load (**0.1x SL**; 0.68 g/L organics), undiluted EPA soil load (**1x SL;** 6.8 g/L organics), mucin component of undiluted soil load (**1x MUC**; 0.8 g/L), BSA component of undiluted soil load (**1x BSA**; 2.5 g/L), yeast extract component from soil load (**1x YE**; 3.5 g/L). Black dotted line represents log_10_CFU/surface inoculated onto the surfaces. Mean value of 5-8 data points is shown. Values at or above the detection limit of at least three colonies counted in all undiluted plated drops (red dotted line) were used for the statistical analysis of CFU counts. Lower values are shown to demonstrate that growth of viable bacteria was detected in at least some replicates.

Minor drying-related decrease of viability was also seen on stainless steel, where the effect towards *E. coli* in low-organic medium (-0.5 log_10_ from 1x soil load, P<0.0001) was not seen in any of the other exposure media with higher organic content indicating that organic content, but not 0.8 g/L mucin alone, protected bacteria from desiccation. The latter disproves the idea that mucin could exclusively protect bacteria against drying-related inactivation similarly to enveloped viruses [35].

## Conclusions

Here we carried out a systematic series of experiments in variable experimental conditions to understand, how relative air humidity of the test environment, droplet size of the bacterial inoculum and organic surface soiling affect antibacterial activity of solid surfaces. Antibacterial activity of five copper, silver and quaternary ammonium-based surfaces was compared against control surfaces of stainless steel and glass using modified ISO 22196 condition which requires bacterial exposure in a thin layer of liquid in nutrient-poor medium, and bacterial exposure in high- or low-organic droplets that were exposed to variable relative air humidity. Across the conditions tested, copper-based surfaces proved to be the most efficient followed by silver and Si-Quat surfaces with the latter showing potential effect specifically towards *S. aureus*. Although several surfaces elicited >3 log_10_ reduction in bacterial viability in different test conditions after 6 h exposure, the regulatory goal of at least 3 log_10_ reduction in up to 1-2 h was only met in case of copper-based surfaces in modified ISO protocol or in low-organic exposures in droplet exposures.

Our results from droplet exposures indicated that antibacterial activity of the tested surfaces against *E. coli* and *S. aureus* decreased with decreasing RH and increasing organic soiling. At low RH, none of the surfaces exhibited biologically or statistically significant antibacterial activity, except for copper in large droplets with low organic content. At high RH, copper surfaces were found to be most efficient in the case of microdroplet inoculation. Results on Si-Quat surfaces showed antibacterial effect only in modified ISO 22196 format and in large low-organic droplets, but high variability of the results made it impossible to draw further conclusions.

The studied environmental variables: humidity and organic content of the medium affected the viability of both *E. coli* and *S. aureus*. As a general rule, on surfaces with lower to no antibacterial activity, viability of *E. coli* was more affected by drying analyzed as a combined effect of RH and inoculum droplet size while *S. aureus* was more affected by the selection of exposure medium. To study the effect of exposure medium on antibacterial effect, we compared the antibacterial activity of copper as the most effective surface and stainless steel as the control surface, in high organics containing soil load and its components compared with low organics environment and showed that the organics in exposure medium efficiently inactivated the antibacterial activity of metal-based surfaces.

There has been a lot of discussion along with a recent ISO initiative to develop a dry standard test method (ISO/DIS 7581) alongside the current wet ISO 22196 industrial standard. However, practical challenges in testing as well as knowledge gaps regarding the effects of environmental variables, especially drying, remain. Our study highlights the need for an open discussion about the testing conditions with an application-relevant test protocol beyond the wet *versus* dry formats and include spray inoculation as one of the worst-case scenarios to accurately assess the antimicrobial activity of solid surfaces in real-life like conditions.

## Data availability

Raw viability data can be found in the Supplementary Raw Data file. Additional information can be obtained from the authors.

## Acknowledgements

This study was funded by the European Union under grant agreement 101057961. Financial support was also received from Estonian Research Council grants PRG1496 and Estonian Centre of Excellence in Research Project “Advanced materials and high-technology devices for sustainable energetics, sensorics and nanoelectronics” TK141 (2014-2020.4.01.150-011). The research was partly conducted using the NAMUR+ core facility funded by projects “Center of nanomaterials technologies and research” (2014-2020.4.01.16-0123) and TT13. This work has been partially supported by Graduate School of Functional materials and technologies receiving funding from the European Regional Development Fund in University of Tartu, Estonia. Featured image icons created by Freepik – Flaticon (www.flaticon.com).

## Author contributions

HR – antibacterial testing, data management, figures preparation, editing of the manuscript.

MR – experimental design of antibacterial testing, data analysis and interpretation, figure design, writing of manuscript.

MK – physico-chemical characterization of surfaces, data interpretation.

DD - physico-chemical characterization of surfaces, data interpretation.

VK – data interpretation, editing of manuscript, project administration.

AI – study design, data interpretation, writing of manuscript, project administration.

