## Supplementary Figures and Tables for "Antibacterial activity of solid surfaces is critically dependent on relative humidity, inoculum volume and organic soiling"

* shared first authorship

| **Abbreviations** | |
|---|---|
| **CuC** | Metallic copper |
| **CuT** | Iniesta CovidSafe copper tape |
| **AgC** | Metallic silver |
| **AgP** | TOUCH Antimicrobial silver paint |
| **SQ** | Si-Quat coating |
| **SS** | Stainless steel |
| **NC** | Borosilicate glass |
| **XPS** | X-ray photoelectron spectroscopy |
| **SEM** | Scanning Electron Microscopy |
| **UHV** | Ultra High Vacuum |
| **RH** | Relative Humidity |
| **NB** | 500-fold diluted nutrient broth |
| **SL** | Soil load |
| **MUC** | Mucin component of undiluted soil load |
| **BSA** | Bovine Serum Albumin component of undiluted soil load |
| **YE** | Yeast extract component from soil load |
| **LB** | Lysogeny broth |
| **SCDLP** | Soya Casein Digest Lecithin Polysorbate Broth |

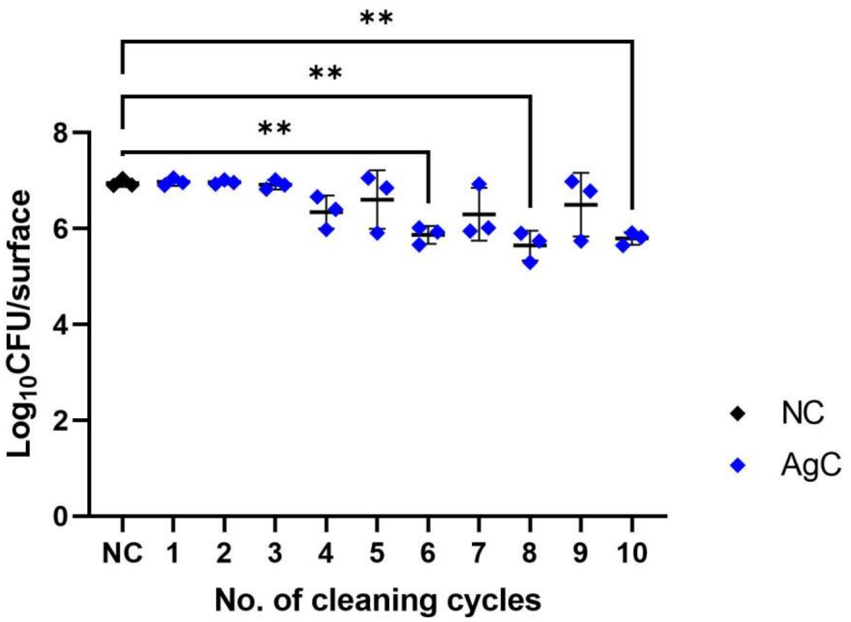


**Figure S1.** Antibacterial activity (1 h exposure in 50 µl soil load medium, 90 % RH) of AgC (metallic silver) surfaces (blue) compared to NC (borosilicate glass) (black) towards *E. coli* after up to 9 additional successive cleaning cycles. A statistically significant linear negative trend from 7.0±0.081 $log_{10}$CFU/surface after 1st cycle to 5.8±0.13 $log_{10}$CFU survivors per surface after 10th cycle was detected after one-way ANOVA analysis (P<0.0001) translating into no statistically significant reduction in viability during the first 5 cleaning cycles and up to 1.2±0.29 logs reduction compared to NC by the last cleaning cycle. The surfaces demonstrate stable activity for the first 3 cleaning cycles and there seems to be a slight increase in antibacterial activity after 5 cleaning cycles. Surfaces were retrieved from the successive cleaning cycles in triplicates and tested for their antibacterial activity in parallel in the same test conditions, with mean and standard deviation presented on the figure. Statistically significant differences from NC are marked with ** (0.01>P>0.001).

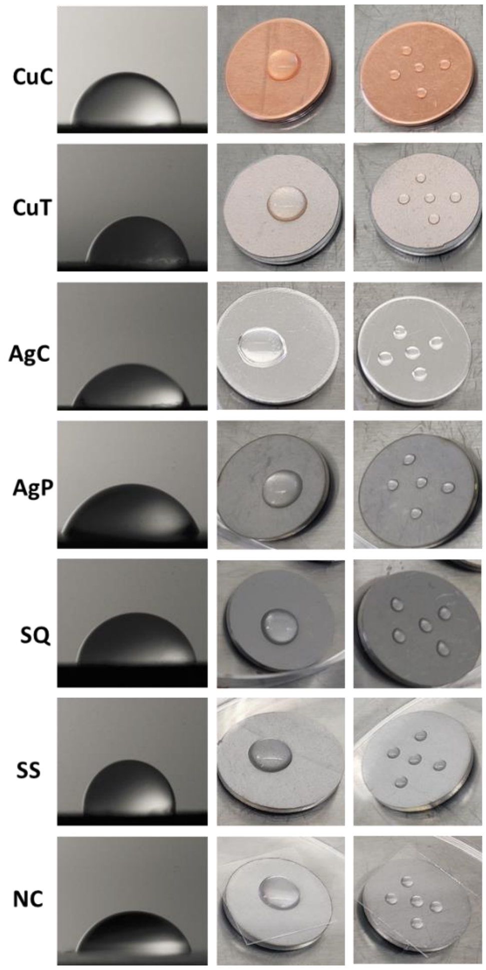


**Figure S2.** Deionized water contact angle on the tested and control surfaces. (CuC, metallic copper, CuT, CovidSafe copper tape, AgC, metallic silver, AgP, TOUCH Antimicrobial silver paint, SQ, Si-Quat coating, SS, stainless steel, NC, borosilicate glass. Left: contact angle, middle: 50 µL drop of bacterial inoculum on the surface, right: 5x2 µL drop of bacterial inoculum on the surface.

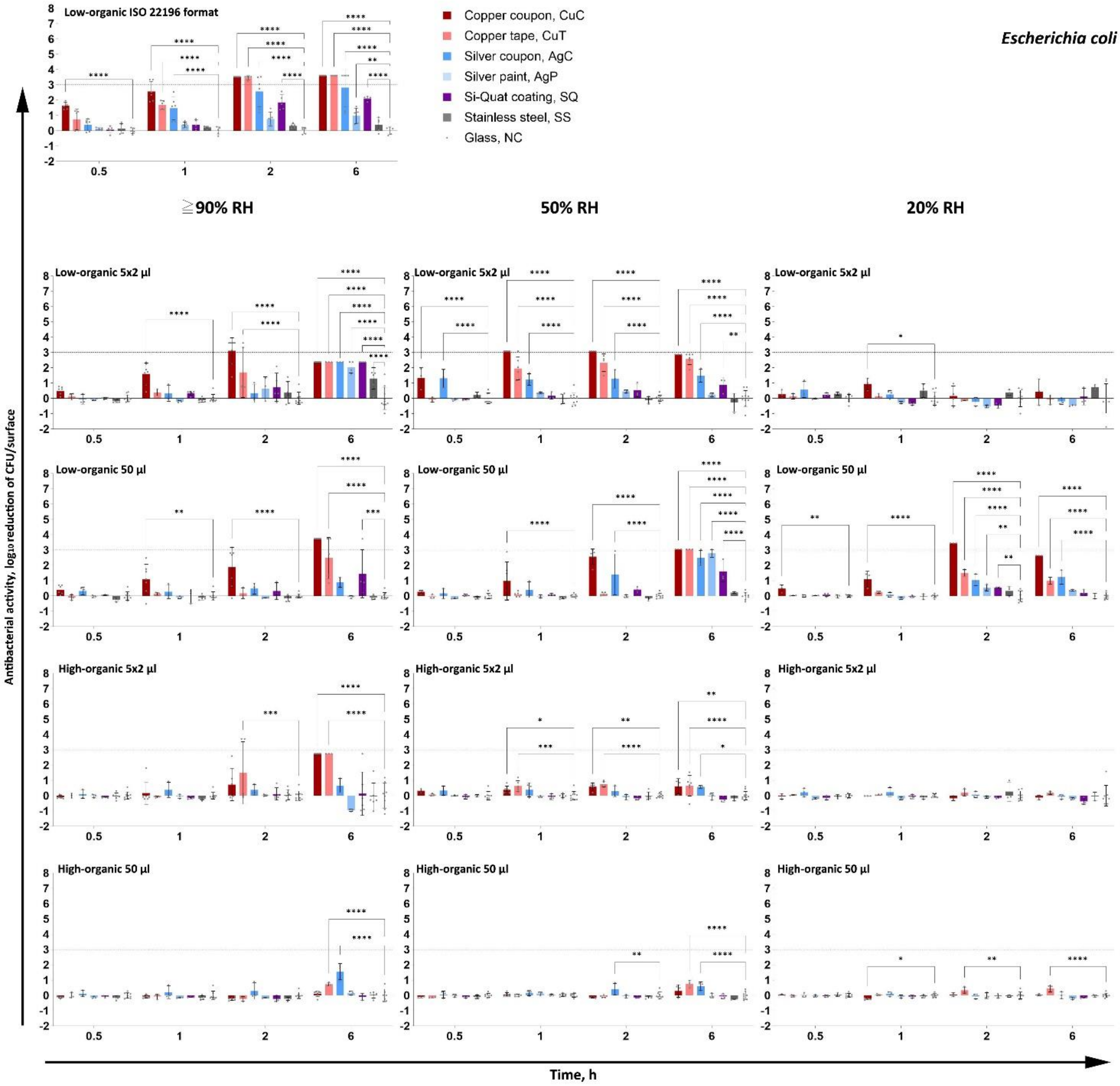


**Figure S3.** Antibacterial activity of the surfaces towards *E. coli* exposure in low organic (A, B, D) and in high organic (C, E) media exposed either as a liquid layer in ISO 22196 format (A), microdroplets (5x2 µl; B, C) or large droplet (50 µl; D, E). $Log_{10}$ reduction is calculated from NC. The maximum possible $log_{10}$ reduction detectable depended on bacterial viability on glass control surface in each time point and test conditions and can be recognized from high antibacterial activity values with no error bars. An average of 3-6 biological replicates with standard deviation is shown. Statistically significant difference (P<0.05) from the NC control surface is marked with **** (P≤ 0.0001), *** (P≤0,001), ** (P≤ 0.01), * (P<0.05). The target of 3 $log_{10}$ reduction is displayed as a grey dotted line.

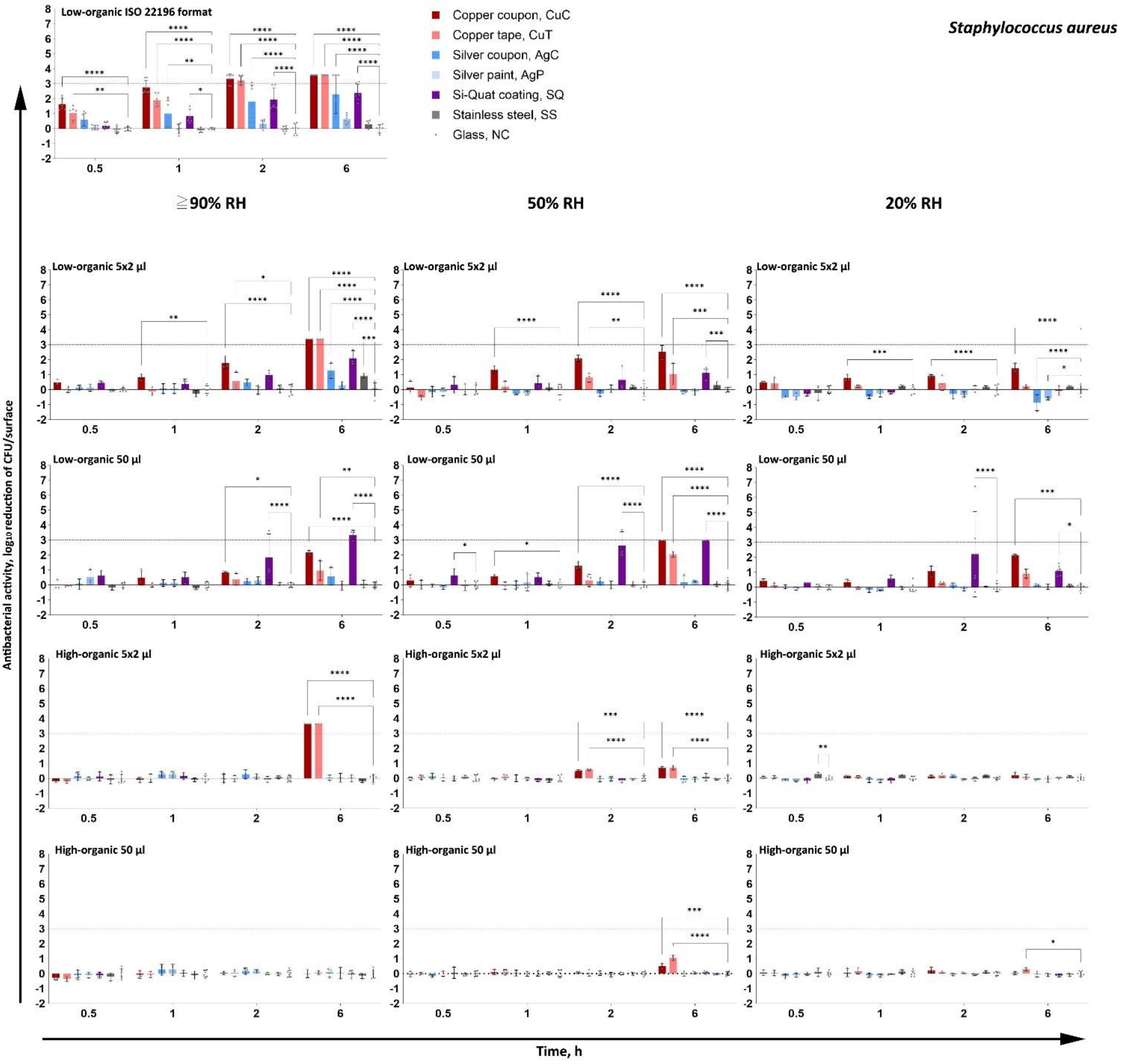


**Figure S4.** Antibacterial activity of the surfaces towards *S. aureus* exposure in low organic (A, B, D) and in high organic (C, E) media exposed either as a liquid layer in ISO 22196 format (A), microdroplets (5x2 µl; B, C) or large droplet (50 µl; D, E). $Log_{10}$ reduction is calculated from NC. The maximum possible $log_{10}$ reduction detectable depended on bacterial viability on glass control surface in each time point and test conditions and can be recognized from high antibacterial activity values with no error bars. An average of 3-6 biological replicates with standard deviation is shown. Statistically significant difference (P<0.05) from the NC control surface is marked with **** (P≤ 0.0001), *** (P≤0,001), ** (P≤ 0.01), * (P<0.05). The target of 3 $log_{10}$ reduction is displayed as a grey dotted line.

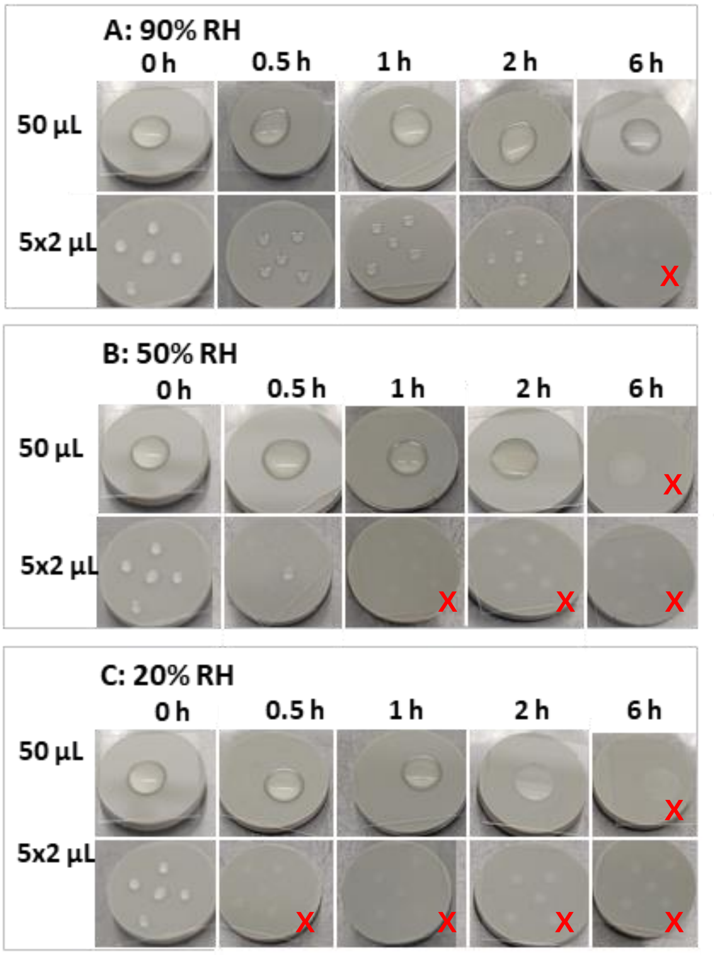


**Figure S5.** Drying of bacterial inoculum on control glass surfaces at different RH in large 50 µL droplets and 2 µL microdroplets. Representative images of droplets at different RH are shown. There were no visible differences in drying of droplets in different media. However, uneven drying time of droplets on the same surface was often observed (middle panel, 0.5 h), possibly explained by spatial variability of the surface characteristics, ventilation in the climate chamber, variation in pipetting or combination of the above. Surfaces with completely dried droplets are marked with a red **X**.

**Table S1.** $Log_{10}$ transformed **viable counts** (in bold) of *E. coli* and *S. aureus* inoculated in droplets on all surfaces in all time-points and *standard deviation* (in italic). Log reduction from initial inoculum is color-coded. Orange color indicates a value below limit of detection (at least 3 colonies counted: 3.18 and 1.48 $log_{10}$ CFU/surface for droplet inoculation and ISO 22196 format respectively).

| *E.coli* | *S.aureus* |
|---|---|
| 0<log rd≤1 | 0<log rd≤1 |
| 1<log rd≤2 | 1<log rd≤2 |
| 2<log rd≤3 | 2<log rd≤3 |
| 3<log rd≤3.8 | 3<log rd≤3.7 |
| log rd>3.82 | log rd>3.67 |
| Mean CFU values below LOD | |

| Species | | *Escherichia coli* | | | | | | | | | | *Staphylococcus aureus* | | | | | | | | | |
|---|---|---|---|---|---|---|---|---|---|---|---|---|---|---|---|---|---|---|---|---|---|
| Exposure medium | | High-organic | | | | | Low-organic | | | | | High-organic | | | | | Low-organic | | | | |
| Time/ Condition | | 0h | 0.5h | 1h | 2h | 6h | 0h | 0.5h | 1h | 2h | 6h | 0h | 0.5h | 1h | 2h | 6h | 0h | 0.5h | 1h | 2h | 6h |
| Copper coupon (CuC) | 90% 50 µl | 7.0 | 7.0 | 6.9 | 7.1 | 7.3 | 7.0 | 6.5 | 5.8 | 5.0 | 0.0 | 6.7 | 6.9 | 6.9 | 6.9 | 7.0 | 6.7 | 6.7 | 6.3 | 6.0 | 4.5 |
| | (SD) | 0.2 | 0.1 | 0.1 | 0.1 | 0.2 | 0.2 | 0.3 | 1.0 | 1.3 | 0.0 | 0.2 | 0.1 | 0.3 | 0.1 | 0.3 | 0.2 | 0.3 | 0.5 | 0.1 | 0.1 |
| | 90% 5x2 µl | 7.0 | 7.1 | 6.7 | 6.2 | 0.0 | 7.0 | 6.4 | 5.3 | 1.7 | 0.0 | 6.7 | 6.9 | 7.0 | 6.9 | 0.0 | 6.7 | 6.2 | 5.7 | 4.8 | 0.0 |
| | (SD) | 0.2 | 0.1 | 0.7 | 1.0 | 0.0 | 0.2 | 0.3 | 0.7 | 2.4 | 0.0 | 0.2 | 0.1 | 0.2 | 0.3 | 0.0 | 0.2 | 0.2 | 0.2 | 0.4 | 0.0 |
| | 50% 50 µl | 7.0 | 7.1 | 7.0 | 7.1 | 6.3 | 7.0 | 6.6 | 6.0 | 4.3 | 0.0 | 6.8 | 7.0 | 6.9 | 6.9 | 6.5 | 6.8 | 6.4 | 6.3 | 5.6 | 0.0 |
| | (SD) | 0.2 | 0.0 | 0.1 | 0.0 | 0.4 | 0.2 | 0.1 | 1.2 | 0.5 | 0.0 | 0.2 | 0.2 | 0.2 | 0.1 | 0.2 | 0.2 | 0.4 | 0.1 | 0.2 | 0.0 |
| | 50% 5x2 µl | 7.0 | 6.6 | 6.4 | 6.2 | 6.0 | 7.0 | 5.5 | 0.0 | 0.0 | 0.0 | 6.9 | 7.0 | 6.9 | 6.3 | 6.2 | 6.9 | 6.4 | 4.9 | 4.1 | 3.4 |
| | (SD) | 0.1 | 0.1 | 0.3 | 0.2 | 0.5 | 0.1 | 0.7 | 0.0 | 0.0 | 0.0 | 0.1 | 0.1 | 0.1 | 0.1 | 0.1 | 0.1 | 0.4 | 0.3 | 0.2 | 0.5 |
| | 20% 50 µl | 7.0 | 7.0 | 7.2 | 6.7 | 6.6 | 7.0 | 6.5 | 5.8 | 0.0 | 0.0 | 7.0 | 7.0 | 6.9 | 6.8 | 6.9 | 7.0 | 6.4 | 6.4 | 5.7 | 4.2 |
| | (SD) | 0.1 | 0.0 | 0.1 | 0.1 | 0.1 | 0.1 | 0.2 | 0.5 | 0.0 | 0.0 | 0.2 | 0.2 | 0.3 | 0.2 | 0.1 | 0.2 | 0.2 | 0.2 | 0.3 | 0.1 |
| | 20% 5x2 µl | 7.1 | 6.8 | 6.7 | 6.8 | 6.4 | 7.1 | 5.8 | 4.7 | 4.9 | 3.8 | 7.0 | 6.9 | 6.8 | 6.8 | 6.8 | 7.0 | 6.0 | 5.6 | 5.4 | 4.7 |
| | (SD) | 0.1 | 0.2 | 0.0 | 0.1 | 0.1 | 0.1 | 0.3 | 0.4 | 0.6 | 1.1 | 0.1 | 0.1 | 0.1 | 0.1 | 0.2 | 0.1 | 0.1 | 0.2 | 0.1 | 0.3 |
| Copper tape (CuT) | 90% 50 µl | 7.0 | 7.0 | 6.9 | 7.1 | 7.3 | 7.0 | 7.0 | 6.7 | 6.7 | 3.2 | 6.7 | 7.0 | 6.9 | 6.9 | 6.9 | 6.7 | 6.7 | 6.8 | 6.4 | 5.7 |
| | (SD) | 0.2 | 0.2 | 0.2 | 0.2 | 0.1 | 0.2 | 0.2 | 0.1 | 0.4 | 3.0 | 0.2 | 0.2 | 0.2 | 0.2 | 0.1 | 0.2 | 0.0 | 0.1 | 0.4 | 0.7 |
| | 90% 5x2 µl | 7.0 | 7.1 | 6.7 | 6.2 | 0.0 | 7.0 | 6.8 | 6.5 | 4.5 | 0.0 | 6.7 | 6.9 | 6.9 | 6.9 | 0.0 | 6.7 | 6.8 | 6.7 | 6.1 | 0.0 |
| | (SD) | 0.2 | 0.2 | 0.1 | 3.1 | 0.0 | 0.2 | 0.2 | 0.2 | 2.8 | 0.0 | 0.2 | 0.1 | 0.3 | 0.2 | 0.0 | 0.2 | 0.2 | 0.3 | 0.6 | 0.0 |
| | 50% 50 µl | 7.0 | 7.1 | 7.0 | 7.1 | 6.3 | 7.0 | 6.9 | 6.8 | 6.8 | 0.0 | 6.8 | 7.0 | 6.9 | 6.9 | 5.9 | 6.8 | 6.7 | 6.9 | 6.6 | 4.1 |
| | (SD) | 0.2 | 0.0 | 0.1 | 0.1 | 0.2 | 0.2 | 0.1 | 0.2 | 0.1 | 0.0 | 0.2 | 0.1 | 0.2 | 0.2 | 0.2 | 0.2 | 0.3 | 0.2 | 0.4 | 0.2 |
| | 50% 5x2 µl | 7.0 | 6.6 | 6.4 | 6.2 | 6.0 | 7.0 | 7.0 | 4.2 | 3.9 | 2.3 | 6.9 | 6.9 | 6.7 | 6.2 | 6.2 | 6.9 | 7.0 | 6.0 | 5.4 | 5.0 |
| | (SD) | 0.1 | 0.1 | 0.3 | 0.2 | 0.7 | 0.1 | 0.2 | 0.9 | 0.6 | 1.8 | 0.1 | 0.2 | 0.2 | 0.1 | 0.1 | 0.1 | 0.1 | 0.3 | 0.3 | 0.7 |
| | 20% 50 µl | 7.0 | 7.0 | 7.2 | 6.7 | 6.6 | 7.0 | 7.0 | 6.6 | 5.1 | 4.8 | 7.0 | 7.0 | 6.8 | 7.0 | 6.7 | 7.0 | 6.7 | 6.8 | 6.4 | 5.4 |
| | (SD) | 0.1 | 0.1 | 0.1 | 0.2 | 0.2 | 0.1 | 0.0 | 0.1 | 0.2 | 0.2 | 0.2 | 0.2 | 0.2 | 0.1 | 0.1 | 0.2 | 0.1 | 0.1 | 0.1 | 0.3 |
| | 20% 5x2 µl | 7.1 | 6.8 | 6.7 | 6.8 | 6.4 | 7.1 | 6.0 | 5.5 | 5.2 | 4.5 | 7.0 | 6.9 | 6.8 | 6.8 | 6.9 | 7.0 | 6.0 | 6.2 | 5.9 | 5.9 |
| | (SD) | 0.1 | 0.0 | 0.0 | 0.2 | 0.1 | 0.1 | 0.2 | 0.2 | 0.0 | 0.3 | 0.1 | 0.1 | 0.1 | 0.1 | 0.2 | 0.1 | 0.4 | 0.1 | 0.5 | 0.1 |
| Silver coupon (AgC) | 90% 50 µl | 7.0 | 6.8 | 6.7 | 6.6 | 5.9 | 7.0 | 6.6 | 6.6 | 6.4 | 6.0 | 6.7 | 6.8 | 6.5 | 6.7 | 6.9 | 6.7 | 6.6 | 6.7 | 6.6 | 6.1 |
| | (SD) | 0.2 | 0.2 | 0.4 | 0.5 | 0.5 | 0.2 | 0.2 | 0.5 | 0.4 | 0.3 | 0.2 | 0.3 | 0.3 | 0.2 | 0.3 | 0.2 | 0.3 | 0.2 | 0.2 | 0.6 |
| | 90% 5x2 µl | 7.0 | 6.8 | 6.5 | 6.5 | 5.3 | 7.0 | 7.0 | 6.6 | 6.5 | 0.9 | 6.7 | 6.6 | 6.6 | 6.6 | 6.8 | 6.7 | 6.6 | 6.5 | 6.2 | 5.3 |
| | (SD) | 0.2 | 0.3 | 0.5 | 0.3 | 0.5 | 0.2 | 0.3 | 0.5 | 0.6 | 1.6 | 0.2 | 0.3 | 0.2 | 0.3 | 0.2 | 0.2 | 0.2 | 0.3 | 0.2 | 0.5 |
| | 50% 50 µl | 7.0 | 6.9 | 6.9 | 6.6 | 6.0 | 7.0 | 6.7 | 6.5 | 5.5 | 3.6 | 6.8 | 7.1 | 7.0 | 6.9 | 7.0 | 6.8 | 6.8 | 6.9 | 6.6 | 6.0 |
| | (SD) | 0.2 | 0.2 | 0.2 | 0.4 | 0.3 | 0.2 | 0.3 | 0.5 | 1.3 | 0.8 | 0.2 | 0.1 | 0.2 | 0.1 | 0.2 | 0.2 | 0.1 | 0.3 | 0.2 | 0.4 |
| | 50% 5x2 µl | 7.0 | 6.6 | 6.4 | 6.4 | 6.1 | 7.0 | 5.6 | 5.0 | 5.0 | 4.6 | 6.9 | 6.8 | 6.8 | 6.9 | 7.0 | 6.9 | 6.7 | 6.5 | 6.5 | 6.2 |
| | (SD) | 0.1 | 0.3 | 0.5 | 0.3 | 0.1 | 0.1 | 0.6 | 0.4 | 0.6 | 0.4 | 0.1 | 0.2 | 0.2 | 0.2 | 0.2 | 0.1 | 0.4 | 0.1 | 0.2 | 0.1 |
| | 20% 50 µl | 7.0 | 7.0 | 6.9 | 6.8 | 6.6 | 7.0 | 7.0 | 6.8 | 5.6 | 4.6 | 7.0 | 7.2 | 7.1 | 7.1 | 7.0 | 7.0 | 6.8 | 6.9 | 6.6 | 6.2 |
| | (SD) | 0.1 | 0.1 | 0.2 | 0.2 | 0.2 | 0.1 | 0.1 | 0.2 | 0.4 | 0.5 | 0.2 | 0.1 | 0.1 | 0.1 | 0.2 | 0.2 | 0.1 | 0.2 | 0.2 | 0.1 |
| | 20% 5x2 µl | 7.1 | 6.5 | 6.4 | 6.5 | 6.4 | 7.1 | 5.5 | 5.4 | 5.3 | 4.6 | 7.0 | 7.1 | 7.0 | 6.8 | 7.1 | 7.0 | 6.9 | 6.9 | 6.6 | 7.0 |
| | (SD) | 0.1 | 0.3 | 0.3 | 0.2 | 0.2 | 0.1 | 0.5 | 0.3 | 0.3 | 0.2 | 0.1 | 0.1 | 0.2 | 0.1 | 0.1 | 0.1 | 0.1 | 0.1 | 0.3 | 0.5 |
| Silver paint (AgP) | 90% 50 µl | 7.0 | 7.0 | 7.0 | 7.0 | 7.3 | 7.0 | 6.9 | 7.0 | 7.0 | 7.0 | 6.7 | 6.8 | 6.5 | 6.7 | 6.9 | 6.7 | 6.2 | 6.7 | 6.5 | 6.7 |
| | (SD) | 0.2 | 0.1 | 0.0 | 0.0 | 0.1 | 0.2 | 0.1 | 0.1 | 0.0 | 0.1 | 0.2 | 0.2 | 0.3 | 0.1 | 0.4 | 0.2 | 0.5 | 0.2 | 0.2 | 0.3 |
| | 90% 5x2 µl | 7.0 | 7.0 | 6.9 | 6.9 | 6.9 | 7.0 | 7.0 | 7.1 | 6.2 | 1.9 | 6.7 | 6.8 | 6.6 | 6.7 | 6.8 | 6.7 | 6.6 | 6.5 | 6.6 | 6.2 |
| | (SD) | 0.2 | 0.1 | 0.1 | 0.1 | 0.1 | 0.2 | 0.1 | 0.1 | 0.8 | 2.2 | 0.2 | 0.1 | 0.2 | 0.2 | 0.3 | 0.2 | 0.2 | 0.3 | 0.3 | 0.2 |
| | 50% 50 µl | 7.0 | 7.0 | 6.9 | 7.0 | 6.6 | 7.0 | 7.0 | 7.0 | 6.9 | 3.4 | 6.8 | 6.9 | 7.0 | 7.1 | 6.9 | 6.8 | 6.9 | 6.7 | 6.9 | 5.9 |
| | (SD) | 0.2 | 0.1 | 0.1 | 0.1 | 0.2 | 0.2 | 0.0 | 0.1 | 0.1 | 0.4 | 0.2 | 0.2 | 0.2 | 0.2 | 0.1 | 0.2 | 0.1 | 0.6 | 0.2 | 0.1 |
| | 50% 5x2 µl | 7.0 | 6.9 | 6.8 | 6.8 | 6.7 | 7.0 | 7.0 | 5.9 | 5.8 | 5.8 | 6.9 | 6.9 | 6.9 | 6.9 | 6.9 | 6.9 | 6.7 | 6.4 | 6.2 | 6.1 |
| | (SD) | 0.1 | 0.0 | 0.1 | 0.2 | 0.2 | 0.1 | 0.1 | 0.1 | 0.1 | 0.1 | 0.1 | 0.2 | 0.1 | 0.1 | 0.2 | 0.1 | 0.3 | 0.1 | 0.2 | 0.2 |
| | 20% 50 µl | 7.0 | 7.1 | 7.0 | 6.8 | 6.8 | 7.0 | 7.0 | 7.1 | 6.1 | 5.5 | 7.0 | 7.1 | 7.1 | 7.0 | 7.0 | 7.0 | 6.9 | 6.9 | 6.8 | 6.3 |
| | (SD) | 0.1 | 0.1 | 0.1 | 0.2 | 0.1 | 0.1 | 0.0 | 0.1 | 0.2 | 0.0 | 0.2 | 0.1 | 0.1 | 0.1 | 0.1 | 0.2 | 0.1 | 0.0 | 0.1 | 0.2 |
| | 20% 5x2 µl | 7.1 | 6.9 | 6.8 | 6.7 | 6.5 | 7.1 | 6.1 | 5.9 | 5.6 | 4.9 | 7.0 | 7.1 | 7.1 | 7.1 | 7.0 | 7.0 | 6.9 | 6.7 | 6.7 | 6.7 |
| | (SD) | 0.1 | 0.1 | 0.1 | 0.1 | 0.1 | 0.1 | 0.0 | 0.1 | 0.1 | 0.1 | 0.1 | 0.1 | 0.1 | 0.1 | 0.2 | 0.1 | 0.2 | 0.3 | 0.2 | 0.1 |

| | Species | *Escherichia coli* | | | | | | | | | | *Staphylococcus aureus* | | | | | | | | | |
|---|---|---|---|---|---|---|---|---|---|---|---|---|---|---|---|---|---|---|---|---|---|
| | Exposure medium | High-organic | | | | | Low-organic | | | | | High-organic | | | | | Low-organic | | | | |
| | Time/ Condition | 0h | 0.5h | 1h | 2h | 6h | 0h | 0.5h | 1h | 2h | 6h | 0h | 0.5h | 1h | 2h | 6h | 0h | 0.5h | 1h | 2h | 6h |
| SiQuat coating (SQ) | 90% 50 µl | **7.0** | **7.0** | **7.0** | **7.1** | **7.5** | **7.0** | **6.8** | **6.9** | **6.6** | **4.7** | **6.7** | **6.8** | **6.8** | **6.9** | **7.0** | **6.7** | **6.1** | **6.3** | **4.8** | **2.1** |
| | | *0.2* | *0.1* | *0.0* | *0.2* | *0.2* | *0.2* | *0.0* | *0.5* | *0.6* | *3.1* | *0.2* | *0.2* | *0.2* | *0.1* | *0.3* | *0.2* | *0.3* | *0.4* | *1.8* | *1.9* |
| | 90% 5x2 µl | **7.0** | **7.1** | **7.0** | **6.8** | **5.8** | **7.0** | **6.9** | **6.6** | **6.1** | **0.0** | **6.7** | **6.6** | **6.7** | **6.9** | **6.8** | **6.7** | **6.3** | **6.2** | **5.6** | **4.4** |
| | | *0.2* | *0.0* | *0.1* | *0.4* | *1.5* | *0.2* | *0.1* | *0.1* | *0.9* | *0.0* | *0.2* | *0.3* | *0.2* | *0.2* | *0.2* | *0.2* | *0.2* | *0.3* | *0.3* | *0.5* |
| | 50% 50 µl | **7.0** | **7.1** | **7.0** | **7.1** | **6.6** | **7.0** | **6.8** | **6.9** | **6.5** | **4.6** | **6.8** | **6.9** | **7.0** | **7.0** | **6.9** | **6.8** | **6.1** | **6.3** | **3.2** | **1.0** |
| | | *0.2* | *0.1* | *0.1* | *0.1* | *0.2* | *0.2* | *0.1* | *0.1* | *0.2* | *0.7* | *0.2* | *0.4* | *0.2* | *0.1* | *0.1* | *0.2* | *0.4* | *0.3* | *2.7* | *1.7* |
| | 50% 5x2 µl | **7.0** | **7.0** | **6.9** | **6.9** | **6.9** | **7.0** | **7.0** | **6.1** | **5.7** | **5.2** | **6.9** | **7.0** | **7.0** | **7.0** | **6.8** | **6.9** | **6.2** | **5.8** | **5.6** | **4.9** |
| | | *0.1* | *0.1* | *0.2* | *0.1* | *0.2* | *0.1* | *0.0* | *0.2* | *0.4* | *0.5* | *0.1* | *0.2* | *0.1* | *0.1* | *0.2* | *0.1* | *0.5* | *0.5* | *0.8* | *0.4* |
| | 20% 50 µl | **7.0** | **7.1** | **7.0** | **6.8** | **6.7** | **7.0** | **6.9** | **6.9** | **6.1** | **5.6** | **7.0** | **7.1** | **7.0** | **7.1** | **7.1** | **7.0** | **6.6** | **6.1** | **4.5** | **5.3** |
| | | *0.1* | *0.1* | *0.1* | *0.1* | *0.0* | *0.1* | *0.1* | *0.1* | *0.1* | *0.2* | *0.2* | *0.1* | *0.0* | *0.1* | *0.1* | *0.2* | *0.0* | *0.2* | *2.9* | *0.3* |
| | 20% 5x2 µl | **7.1** | **6.8** | **6.7** | **6.7** | **6.7** | **7.1** | **5.9** | **5.9** | **5.5** | **4.3** | **7.0** | **7.1** | **7.1** | **7.0** | **6.9** | **7.0** | **6.7** | **6.6** | **6.3** | **6.2** |
| | | *0.1* | *0.1* | *0.1* | *0.1* | *0.2* | *0.1* | *0.1* | *0.1* | *0.2* | *0.5* | *0.1* | *0.2* | *0.2* | *0.1* | *0.1* | *0.1* | *0.2* | *0.1* | *0.2* | *0.3* |
| Stainless steel (SS) | 90% 50 µl | **7.0** | **7.0** | **7.0** | **7.1** | **7.4** | **7.0** | **7.1** | **7.0** | **7.0** | **7.0** | **6.7** | **6.9** | **6.9** | **6.9** | **7.1** | **6.7** | **6.8** | **7.0** | **6.8** | **6.6** |
| | | *0.2* | *0.1* | *0.1* | *0.1* | *0.2* | *0.2* | *0.1* | *0.2* | *0.2* | *0.1* | *0.2* | *0.3* | *0.3* | *0.1* | *0.2* | *0.2* | *0.2* | *0.1* | *0.1* | *0.2* |
| | 90% 5x2 µl | **7.0** | **7.0** | **7.1** | **6.8** | **6.0** | **7.0** | **7.1** | **7.0** | **6.4** | **4.2** | **6.7** | **6.8** | **7.0** | **6.8** | **7.0** | **6.7** | **6.8** | **6.8** | **6.6** | **5.6** |
| | | *0.2* | *0.2* | *0.1* | *0.3* | *0.9* | *0.2* | *0.1* | *0.1* | *0.7* | *0.7* | *0.2* | *0.3* | *0.1* | *0.1* | *0.1* | *0.2* | *0.1* | *0.2* | *0.2* | *0.2* |
| | 50% 50 µl | **7.0** | **7.1** | **7.0** | **7.1** | **6.8** | **7.0** | **7.0** | **7.1** | **7.1** | **6.0** | **6.8** | **7.0** | **7.0** | **7.0** | **7.0** | **6.8** | **6.8** | **6.8** | **6.8** | **6.1** |
| | | *0.2* | *0.1* | *0.1* | *0.1* | *0.0* | *0.2* | *0.1* | *0.1* | *0.1* | *0.1* | *0.2* | *0.1* | *0.1* | *0.1* | *0.1* | *0.2* | *0.3* | *0.2* | *0.2* | *0.2* |
| | 50% 5x2 µl | **7.0** | **6.9** | **6.8** | **6.8** | **6.8** | **7.0** | **6.7** | **6.3** | **6.4** | **6.3** | **6.9** | **6.9** | **7.0** | **6.9** | **6.9** | **6.9** | **6.5** | **6.0** | **6.1** | **5.7** |
| | | *0.1* | *0.1* | *0.1* | *0.3* | *0.2* | *0.1* | *0.2* | *0.3* | *0.2* | *0.6* | *0.1* | *0.1* | *0.1* | *0.1* | *0.1* | *0.1* | *0.3* | *0.2* | *0.1* | *0.2* |
| | 20% 50 µl | **7.0** | **7.0** | **7.0** | **6.8** | **6.6** | **7.0** | **7.0** | **6.9** | **6.3** | **5.8** | **7.0** | **6.9** | **6.9** | **6.9** | **7.0** | **7.0** | **6.8** | **6.8** | **6.7** | **6.2** |
| | | *0.1* | *0.1* | *0.1* | *0.1* | *0.1* | *0.1* | *0.1* | *0.2* | *0.3* | *0.2* | *0.2* | *0.3* | *0.2* | *0.1* | *0.1* | *0.2* | *0.2* | *0.1* | *0.0* | *0.1* |
| | 20% 5x2 µl | **7.1** | **6.8** | **6.7** | **6.3** | **6.3** | **7.1** | **5.8** | **5.1** | **4.7** | **3.7** | **7.0** | **6.7** | **6.7** | **6.8** | **6.8** | **7.0** | **6.6** | **6.2** | **6.1** | **5.9** |
| | | *0.1* | *0.1* | *0.1* | *0.6* | *0.3* | *0.1* | *0.1* | *0.5* | *0.2* | *0.2* | *0.1* | *0.1* | *0.0* | *0.1* | *0.1* | *0.1* | *0.5* | *0.1* | *0.1* | *0.1* |
| Negative control (NC) | 90% 50 µl | **7.0** | **6.9** | **6.9** | **6.9** | **7.4** | **7.0** | **6.9** | **6.9** | **6.9** | **6.9** | **6.7** | **6.7** | **6.8** | **6.9** | **7.0** | **6.7** | **6.7** | **6.8** | **6.8** | **6.7** |
| | | *0.2* | *0.2* | *0.3* | *0.2* | *0.4* | *0.2* | *0.3* | *0.3* | *0.1* | *0.2* | *0.2* | *0.3* | *0.2* | *0.2* | *0.3* | *0.2* | *0.2* | *0.2* | *0.1* | *0.2* |
| | 90% 5x2 µl | **7.0** | **7.0** | **6.9** | **6.9** | **5.9** | **7.0** | **6.9** | **6.9** | **6.8** | **5.6** | **6.7** | **6.7** | **6.9** | **6.9** | **6.8** | **6.7** | **6.7** | **6.6** | **6.6** | **6.5** |
| | | *0.2* | *0.2* | *0.2* | *0.3* | *0.8* | *0.2* | *0.2* | *0.3* | *0.4* | *0.7* | *0.2* | *0.3* | *0.2* | *0.2* | *0.2* | *0.2* | *0.1* | *0.2* | *0.4* | *0.5* |
| | 50% 50 µl | **7.0** | **7.0** | **7.1** | **7.0** | **6.6** | **7.0** | **6.9** | **6.9** | **6.9** | **6.2** | **6.8** | **6.9** | **7.0** | **7.0** | **7.0** | **6.8** | **6.8** | **6.8** | **6.9** | **6.2** |
| | | *0.2* | *0.1* | *0.1* | *0.2* | *0.3* | *0.2* | *0.2* | *0.1* | *0.2* | *0.2* | *0.2* | *0.2* | *0.2* | *0.1* | *0.1* | *0.2* | *0.2* | *0.2* | *0.2* | *0.3* |
| | 50% 5x2 µl | **7.0** | **6.9** | **6.8** | **6.7** | **6.6** | **7.0** | **6.9** | **6.3** | **6.3** | **6.0** | **6.9** | **6.9** | **6.9** | **6.8** | **6.9** | **6.9** | **6.5** | **6.2** | **6.2** | **6.0** |
| | | *0.1* | *0.3* | *0.3* | *0.1* | *0.3* | *0.1* | *0.3* | *0.4* | *0.2* | *0.5* | *0.1* | *0.2* | *0.2* | *0.2* | *0.2* | *0.1* | *0.3* | *0.3* | *0.4* | *0.1* |
| | 20% 50 µl | **7.0** | **7.0** | **7.0** | **6.8** | **6.6** | **7.0** | **7.0** | **6.9** | **6.6** | **5.8** | **7.0** | **7.0** | **7.0** | **7.0** | **7.0** | **7.0** | **6.8** | **6.7** | **6.7** | **6.3** |
| | | *0.1* | *0.1* | *0.1* | *0.2* | *0.1* | *0.1* | *0.1* | *0.1* | *0.4* | *0.2* | *0.2* | *0.2* | *0.2* | *0.1* | *0.2* | *0.2* | *0.1* | *0.3* | *0.3* | *0.2* |
| | 20% 5x2 µl | **7.1** | **6.7** | **6.6** | **6.6** | **6.3** | **7.1** | **6.1** | **5.6** | **5.1** | **4.4** | **7.0** | **6.9** | **6.9** | **7.0** | **7.0** | **7.0** | **6.4** | **6.4** | **6.3** | **6.1** |
| | | *0.1* | *0.1* | *0.1* | *0.3* | *0.7* | *0.1* | *0.3* | *0.4* | *0.5* | *1.1* | *0.1* | *0.1* | *0.1* | *0.1* | *0.2* | *0.1* | *0.2* | *0.3* | *0.3* | *0.3* |

**Table S2.** $Log_{10}$ transformed **viable counts** (in bold) of *E. coli* and *S. aureus* inoculated in ISO 22196 format on all surfaces in all time-points and *standard deviation* (in italic). Log reduction from initial inoculum is color-coded.

| | *Escherichia coli* | | | | | *Staphylococcus aureus* | | | | |
|---|---|---|---|---|---|---|---|---|---|---|
| **Time** | 0h | 0.5h | 1h | 2h | 6h | 0h | 0.5h | 1h | 2h | 6h |
| **CuC** | **4.7** | **3.0** | **2.1** | **0.3** | **0.3** | **4.7** | **3.3** | **1.7** | **1.3** | **0.3** |
| | *0.1* | *0.2* | *0.9* | *0.5* | *0.5* | *0.1* | *0.4* | *1.2* | *0.9* | *0.5* |
| **CuT** | **4.7** | **3.9** | **3.1** | **0.6** | **0.4** | **4.7** | **3.9** | **3.0** | **1.7** | **0.3** |
| | *0.1* | *0.7* | *0.3* | *0.9* | *0.5* | *0.1* | *0.5* | *0.4* | *0.5* | *0.5* |
| **AgC** | **4.7** | **4.3** | **3.4** | **2.4** | **1.5** | **4.7** | **4.3** | **3.9** | **3.2** | **2.4** |
| | *0.1* | *0.4* | *0.8* | *1.0* | *1.9* | *0.1* | *0.4* | *0.9* | *1.1* | *1.7* |
| **AgP** | **4.7** | **4.5** | **4.5** | **4.2** | **4.1** | **4.7** | **4.8** | **4.9** | **4.7** | **4.4** |
| | *0.1* | *0.1* | *0.2* | *0.5* | *0.5* | *0.1* | *0.1* | *0.3* | *0.3* | *0.3* |
| **SQ** | **4.7** | **4.6** | **4.4** | **3.2** | **3.0** | **4.7** | **4.7** | **4.1** | **3.1** | **2.7** |
| | *0.1* | *0.2* | *0.3* | *0.5* | *0.1* | *0.1* | *0.2* | *0.5* | *0.8* | *0.6* |
| **SS** | **4.7** | **4.5** | **4.6** | **4.7** | **4.7** | **4.7** | **5.0** | **5.0** | **5.1** | **4.8** |
| | *0.1* | *0.3* | *0.1* | *0.1* | *0.5* | *0.1* | *0.2* | *0.1* | *0.2* | *0.2* |
| **NC** | **4.7** | **4.6** | **4.8** | **5.0** | **5.1** | **4.7** | **4.9** | **4.9** | **5.1** | **5.1** |
| | *0.1* | *0.2* | *0.3* | *0.2* | *0.2* | *0.1* | *0.1* | *0.1* | *0.3* | *0.3* |

| ***E.coli*** | ***S.aureus*** |
|---|---|
| 0<log rd≤1 | 0<log rd≤1 |
| 1<log rd≤2 | 1<log rd≤2 |
| 2<log rd≤3 | 2<log rd≤3 |
| log rd>3 | log rd>3 |

**Table S3**. Contribution of **surface type** (*vs* glass control, NC), **exposure medium** (high vs low organic content), **drying of the inoculum** and their interactions (indicated by variable x variable interaction in the legend) to the total variation in viable counts based on 3-way ANOVA analysis of 2 h exposure data of *E. coli* (left) and *S. aureus* (right). The parameters of relative humidity (RH) and inoculum droplet size proved to be interdependent in their effect on viability probably by affecting drying time and could thereby impair main effect and interaction interpretation in multifactorial ANOVA. Instead, a combined variable, drying, was used. As a categorical variable drying comprised of all 6 possible combinations of RH (20%, 50% and 90%) and inoculum droplet sizes (5x2 µl and 50 µl). Green color indicates statistical significance ($P<0.05$), grey color indicates no statistically significant variation ($P\geq0.05$). (CuC – metallic copper; CuT – CovidSafe copper tape; AgC – metallic silver; AgP – TOUCH Antimicrobial silver paint; SQ - Si-Quat coating). Data forming the basis of the analysis is also presented in Supplementary Table S1 (2 h time-point).

| Source of Variation | **% of total variation of viable counts** explained by the environmental variables and interactions of variables based on 3-way ANOVA of 2h data. | | | | | | | | | | | |
|---|---|---|---|---|---|---|---|---|---|---|---|---|
| | ***Escherichia coli*** | | | | | | ***Staphylococcus aureus*** | | | | | |
| | **CuC** | **CuT** | **AgC** | **AgP** | **SQ** | **SS** | **CuC** | **CuT** | **AgC** | **AgP** | **SQ** | **SS** |
| **Drying** (all possible combinations of RH and inoculum droplet size) | 6.25 | 24.52 | 15.59 | 26.60 | 19.72 | 41.58 | 12.66 | 23.86 | 11.92 | 15.91 | 1.37 | 21.18 |
| **Surface type** (vs glass (NC)) | 24.14 | 12.34 | 10.23 | 0.18 | 0.99 | 0.18 | 27.19 | 11.71 | 2.03 | 0.01 | 10.73 | 1.06 |
| **Exposure medium** (low-organic vs high-organic) | 33.82 | 10.82 | 17.35 | 14.31 | 20.19 | 11.71 | 40.64 | 27.92 | 23.93 | 21.47 | 25.88 | 25.65 |
| **Surface type x Drying** interaction | 6.70 | 13.62 | 4.79 | 3.18 | 2.93 | 2.00 | 3.95 | 3.88 | 6.71 | 4.21 | 5.11 | 0.67 |
| **Drying x Exposure medium** interaction | 1.20 | 6.68 | 9.33 | 11.14 | 7.79 | 14.01 | 4.12 | 5.09 | 3.21 | 8.36 | 2.06 | 12.13 |
| **Surface type x Exposure medium** interaction | 18.76 | 1.82 | 2.65 | 0.71 | 2.67 | 0.22 | 16.88 | 2.96 | 0.01 | 0.06 | 12.33 | 0.09 |
| **Drying x Exposure medium x Surface type** interaction | 2.74 | 2.74 | 3.41 | 2.19 | 1.68 | 0.47 | 1.96 | 0.61 | 3.55 | 1.19 | 4.58 | 0.49 |
